# KCa3.1 Contributes to Neuroinflammation and Nigral Dopaminergic Neurodegeneration in Experimental models of Parkinson’s Disease

**DOI:** 10.1101/2025.03.18.643982

**Authors:** Manikandan Samidurai, Karthick Chennakesavan, Souvarish Sarkar, Emir Malovic, Hai M. Nguyen, Latika Singh, Anil Kumar, Alyssa Ealy, Chelva Janarthanam, Bharathi Niveditha Palanisamy, Naveen Kondru, Gary Zenitsky, Huajun Jin, Vellareddy Anantharam, Anumantha Kanthasamy, Hui zhang, Heike Wulff, Arthi Kanthasamy

**Affiliations:** Isakson Center for Neurological Disease Research, Department of Physiology and Pharmacology, University of Georgia, 500 DW Brooks Drive, Athens, GA 30602, USA; Parkinson’s Disorder Research Program, Iowa Center for Advanced Neurotoxicology, Department of Biomedical Sciences, Iowa State University, Ames, IA 50011, USA; Department of Pharmacology, School of Medicine, University of California, Davis, Davis, CA, United States

**Keywords:** α-synuclein, KCa3.1, neuroinflammation, Fyn, STAT1, Parkinson’s disease

## Abstract

Chronic neuroinflammation and misfolded α-synuclein (αSyn) have been identified as key pathological correlates driving Parkinson’s disease (PD) pathogenesis; however, the contribution of ion channels to microglia activation in the context of α-synucleinopathy remains elusive. Herein, we show that KCa3.1, a calcium-activated potassium channel, is robustly upregulated within microglia in multiple preclinical models of PD and, most importantly, in human PD and dementia with Lewy bodies (DLB) brains. Pharmacological inhibition of KCa3.1 via senicapoc or TRAM-34 inhibits KCa3.1 channel activity and the associated reactive microglial phenotype in response to aggregated αSyn, as well as ameliorates of PD like pathology in diverse PD mouse models. Additionally, proteomic and transcriptomic profiling of microglia revealed that senicapoc ameliorates aggregated αSyn-induced, inflammation-associated pathways and dysregulated metabolism in primary microglial cells. Mechanistically, FYN kinase in a STAT1 dependent manner regulates KCa3.1 mediated the microglial reactive activation phenotype after α-synucleinopathy. Moreover, reduced neuroinflammation and subsequent PD-like neuropathology were observed in SYN AAV inoculated KCa3.1 knockout mice. Together, these findings suggest that KCa3.1 inhibition represents a novel therapeutic strategy for treating patients with PD and related α-synucleinopathies.

## Introduction

Parkinson’s disease (PD), the second most common neurodegenerative disorder, constitutes a large unmet medical need and exerts a huge socioeconomic burden on society. Therapeutic drug development strategies have been challenging and several of the approved drugs offer only symptomatic management without treating the underlying causes [1]. PD pathogenesis is complex and is characterized by progressive loss of nigral dopamin(DA)ergic neurons, along with an accompanying accumulation of unfolded aggregated α-synuclein (αSyn_Agg_) within Lewy bodies and Lewy neurites [2, 3]. The loss of extra striatal neurons and non-motor symptoms have also been implicated in the pathogenesis of PD [4, 5]. In recent years, a growing body of clinical evidence and studies conducted in experimental models of PD support a pivotal role for neuroinflammation in PD and dementia with Lewy bodies (DLB) [6–8]. Abnormal forms of αSyn have been thought to propagate in the brain via a prion-like mechanism [9, 10]. Although αSyn has been shown to be abundantly expressed within the neuron, the brain’s resident macrophages, microglia, facilitate αSyn propagation via their ability to promote αSyn uptake [11]. In this context, misfolded αSyn released in the vicinity of dying neurons can elicit detrimental effects on neuronal and glial cells, via seeding of αSyn_Agg_, thereby facilitating disease progression [12–15]. The mechanisms underlying microglia-mediated nigral DAergic neurodegeneration have been the subject of intense investigations. For example, microglial mitochondrial dysfunction, neuroinflammation, lysosomal dysfunction, and alteration of calcium homeostasis have been linked to nigral DAergic neurodegeneration [16, 17]. We have previously shown that αSyn_Agg_ induces a microglial activation response, resulting in nigral DAergic neurodegeneration in a PKCδ-dependent manner [18]. Both TLR2 [19, 20] and CD36 [21] are expressed on microglial cells and have been shown to be involved in the αSyn_Agg_-induced microglial activation response, while inhibition of TLR2 via a neutralizing antibody reduces neuroinflammation and tyrosine hydroxylase (TH^+^) cell loss in an α-synucleinopathy mouse model [22].

KCa3.1 is an intermediate-conductance Ca^2+^-activated K^+^ channel that is expressed in microglia, T-lymphocytes, and macrophages [23–26]. KCa3.1 has been shown to regulate not only cellular migration but also proliferation via K^+^ efflux, which enables the maintenance of a negative membrane potential, thereby facilitating Ca^2+^ influx for various cellular activities [26]. KCa3.1 upregulation has been evidenced in pathological conditions such as AD [26], cerebral ischemia [27], and perioperative cognitive decline [28]. Recently, we reported that the upregulation of KCa3.1 closely parallels microglia-mediated neuroinflammation, amyloid-beta aggregation and hippocampal toxicity in an AD mouse model, suggesting that KCa3.1 exhibits a pathological role in proteinopathies including AD [29, 30]. Furthermore, KCa3.1 blockers have been shown to ameliorate the reactive microglia phenotype, amyloid-beta burden, and hippocampal neurotoxicity [30]. However, whether KCa3.1 influences neuroinflammation and nigral DAergic neurodegeneration in the context of α-synucleinopathy remains unknown.

In this study, we show that pharmacological blockade of KCa3.1 ameliorated αSyn_Agg_-induced neuroinflammation both *in vitro* and *in vivo*, and preserved nigral DAergic neuronal integrity. Our mechanistic studies revealed the central role of the Fyn/STAT1 proinflammatory signaling axis in regulating both the microglial activation response and KCa3.1 expression after α-synucleinopathy. Moreover, our studies revealed that KCa3.1 knockout ameliorated neuroinflammation, αSyn pathology, and nigral DAergic neurodegeneration in an α-synucleinopathy mouse model. Together, our data from diverse experimental models of PD suggest that KCa3.1 plays a pivotal role in mediating neuroinflammation and associated nigral DAergic neurodegeneration in experimental Parkinsonism, highlighting the potential utility of KCa3.1 as a therapeutic target for PD related synucleinopathies.

## RESULTS

### KCa3.1 expression is increased in the brains of PD and DLB patients and preclinical mouse models of PD

To test whether chronic inflammatory conditions during PD alter KCa3.1 expression, we determined the KCa3.1 expression levels in the nigral brain region of postmortem PD and DLB brains and in age-matched controls. Western blot (WB) analysis reveals increased KCa3.1 expression in the midbrain region of postmortem PD brains as compared to age-matched controls (Fig. 1A). Upon further characterization of KCa3.1 expression, our IHC studies reveal increased localization of KCa3.1 within reactive amoeboid microglia in PD and DLB postmortem brains as compared to control patients (Fig. 1B-C). We further confirmed this finding in MPTP-treated mice whereby the expression of KCa3.1 was elevated, as demonstrated by both WB (Fig. 1D) and KCa3.1/IBA-1 double immunolabeling studies in the substantia nigra (Fig. 1E). Our finding is in line with a previous finding showing that KCa3.1 was upregulated in both AD postmortem brains and in 5xFAD mice [30]. Next, we examined KCa3.1 expression in MitoPark (MP) mice, which exhibit a heightened microglial activation response in close association with progressive loss of nigral DAergic processes and motor deficits over a 32-wk period [31, 32]. Our WB studies reveal a significantly increased expression of KCa3.1 in the nigra of MP mice as compared to littermate controls at 24 wk (Fig. 1F). Notably, KCa3.1 expression was confined within reactive amoeboid microglia in the nigra of MP mice (Fig. 1G), which closely paralleled elevated levels of KCa3.1 mRNA expression in MP mice compared to littermate controls (Fig. 1H). Taken together, these results suggest a role for microglial KCa3.1 during neuroinflammation in PD.

**Fig 1.**
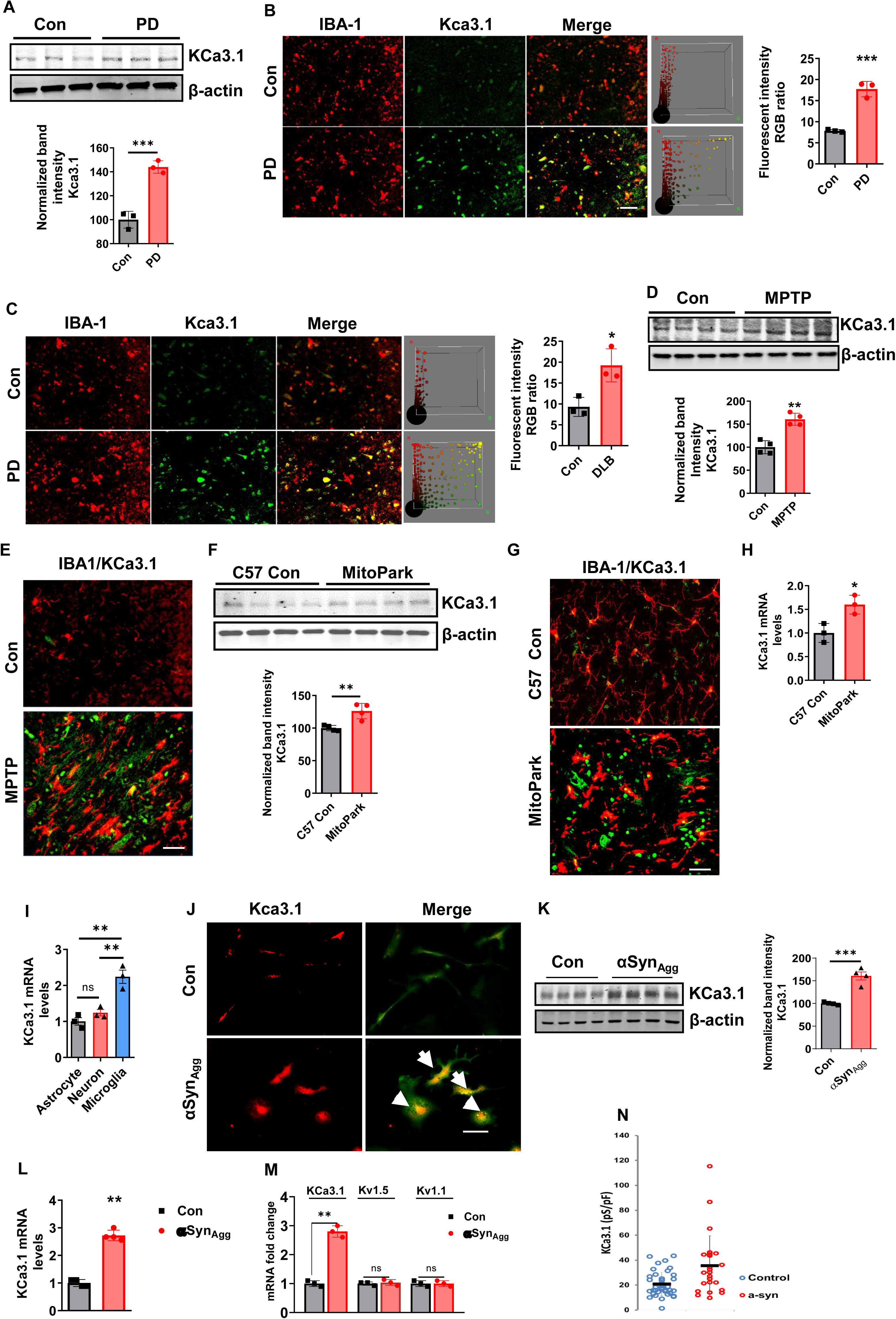
Increased expression of KCa3.1 within reactive microglia in post-mortem PD brains, mouse models of PD and αSyn_Agg_-stimulated mouse primary microglial cells. **(A)** Immunoblotting analysis and densitometric quantification of KCa3.1 normalized to β-actin, an internal loading control. n=3/group; unpaired two-tailed t-test. **(B)** Representative immunostaining images and quantification of KCa3.1/IBA-1 colocalization in the nigra of PD and control brains. Scale bar, 10 µM. Data represents the average of 3 Con and 3 PD brains. **(C)** Representative immunostaining of KCa3.1 in slices from the substantia nigra from control and DLB patients showing colocalization between KCa3.1 (green) and the microglial marker HLA-DR (red). Scale bar, 10 µM. Right: quantification of KCa3.1 immunofluorescence intensity; n=3 from Con and DLB brains. **(D)** Immunoblot analysis of KCa3.1 expression in the ventral midbrain region of MPTP- and saline-treated mice, conducted at 14 d after MPTP injection, and densitometric scanning analysis of KCa3.1 levels (n=4 mice per group). **(E)** Representative dual immunolabeling images for KCa3.1/IBA-1 in the ventral midbrain region following MPTP treatment (n=3 mice per group). **(F)** Western blot (WB) analysis for KCa3.1 in the substantia nigra tissue from MitoPark mice (MP) and littermate controls (C57 Con) at 24 wk of age. Bottom, densitometric scanning analysis of KCa3.1 levels (n=4 mice per group). **(G)** Co-staining with microglial marker IBA-1 in the nigra of MP mice revealed that KCa3.1 is highly expressed within IBA-1-positive amoeboid microglial cells. Images were acquired from 24-wk-old littermate control and MP mice. Scale bar is 100 µM. **(H)** qRT-PCR analysis for KCa3.1 in MP mice at 24 wk of age (n=3). **(I)** RT-qPCR analysis using mRNA generated from cultured primary mouse microglia, astrocytes, and neurons showing that KCa3.1 is highly expressed in microglia. **(J)** KCa3.1 expression was localized to reactive microglia (amoeboid shaped) in αSyn_Agg_-stimulated primary microglia cells. **(K)** Validation of increased expression of KCa3.1 in αSyn_Agg_-stimulated primary microglia by WB analysis. Right: Quantification of KCa3.1 expression by densitometric scanning analysis with β-actin used as an internal control. **(L)** RT-qPCR analysis showing transcriptional upregulation of KCa3.1 in primary murine microglial cultures stimulated with αSyn_Agg_ for 24 h. **(M)** RT-qPCR analysis revealing increased mRNA expression of KCa3.1 without alteration of potassium channels Kv1.5 or Kv1.1 in αSyn_Agg_-stimulated primary microglia. **(N)** Scatter plot displaying greater KCa3.1 channel current density in αSyn_Agg_-treated compared to untreated primary microglia. Data are mean ± SEM with 3-6 biological replicates from 3-4 independent experiments unless otherwise noted. Student’s t-tests or one-way ANOVA with Bonferroni multiple comparison tests were utilized for statistical analysis. *p≤0.05, **p<0.01, *** p<0.001; ns, not significant.

### Functional characterization of KCa3.1 expression in primary microglial cells stimulated with aggregated αSyn

To characterize the αSyn_Agg_-induced immune responses, we initially confirmed the size of fibrils using transmission electron microscopy (TEM) [33] (Fig. S1). Our RT-qPCR analysis reveals increased expression of KCa3.1 in primary microglial cells, as compared to astrocytes and neural cells (Fig. 1I). To further investigate whether αSyn_Agg_ promotes KCa3.1 activation, we established mixed glial cultures and subsequently treated the isolated primary microglia with 1 µM αSyn_Agg_ for 24-48 h. To identify the effect of αSyn_Agg_ on KCa3.1 expression, we performed KCa3.1/IBA-1 double immunolabeling studies, which revealed elevated KCa3.1 expression within amoeboid primary microglia after αSyn_Agg_ treatment as compared to control cells (Fig. 1J). Parallel WB (Fig. 1K) and RT-qPCR (Fig. 1L) analyses reveal increased protein and mRNA expression of KCa3.1 in αSyn_Agg_-stimulated primary microglial cells. Additionally, we demonstrated that αSyn_Agg_ significantly upregulated KCa3.1 mRNA expression as compared with Kv1.1, or Kv1.5 (Fig. 1M) [34], confirming the specificity of KCa3.1 after αSyn_Agg_ stimulation. Our findings are in accordance with Qin et al.’s [35] RNA-seq finding demonstrating an 8-fold increase in KCNN4 (gene encoding KCa3.1) expression in the midbrain regions of rAAV-αSyn-inoculated rats as compared to empty vector-inoculated animals. Similarly, whole-cell patch-clamping revealed that αSyn_Agg_ treatment increased functional KCa3.1 channel activity (Fig. 1N), suggesting that KCa3.1 expression may regulate microglia-mediated inflammatory response. Collectively, these findings demonstrate that inflammatory stimuli such as αSyn_Agg_ induce functionally relevant increases in microglial KCa3.1 protein and mRNA levels, which may partly explain the increased KCa3.1 expression evidenced in PD brains.

We next determined whether KCa3.1 activation contributes to αSyn_Agg_-induced upregulation of proinflammatory cytokines. The αSyn_Agg_-induced KCa3.1 current can be blocked by the KCa3.1 inhibitor TRAM-34 in a whole-cell patch-clamp experiment (Fig. 2A). To determine whether TRAM-34 ameliorates the αSyn_Agg_-induced reactive microglial phenotype in primary murine microglial cultures, we treated primary murine microglia isolated from neonatal mice with 1 µM αSyn_Agg_ in the presence or absence of 1 µM of TRAM-34 for 48 h. TRAM-34 treatment markedly reduced the release of proinflammatory cytokines, including IL-1β, IL-6, TNF-α, and IL-12 (Fig. 2B-E). We further corroborated these findings by employing another clinically tested KCa3.1 blocker, senicapoc (Fig. S1D). In primary cultured microglia co-treated with 1 μM αSyn_Agg_ and 1 µM of senicapoc for 17 h, we observed a marked decrease in the mRNA expression of proinflammatory molecules TNFα, IL-1β, and iNOS (Fig. 2F-H). Additionally, senicapoc co-treatment significantly inhibited αSyn_Agg_-induced nitrite release (Fig. 2I). Next, we performed an siRNA-mediated KCa3.1 knockdown experiment in primary murine microglial cultures. ELISA analysis revealed a significant reduction of αSyn_Agg_-induced proinflammatory IL-1β release in the cell supernatants of KCa3.1-siRNA-transfected cells, compared to non-specific siRNA-treated cells (Fig. 2J). Collectively, these results indicate that KCa3.1 regulates reactive microgliosis and that pharmacological inhibition of KCa3.1 may serve as a viable therapeutic strategy for the treatment of α-synucleinopathies and other neuroinflammation-related disorders.

**Fig 2.**
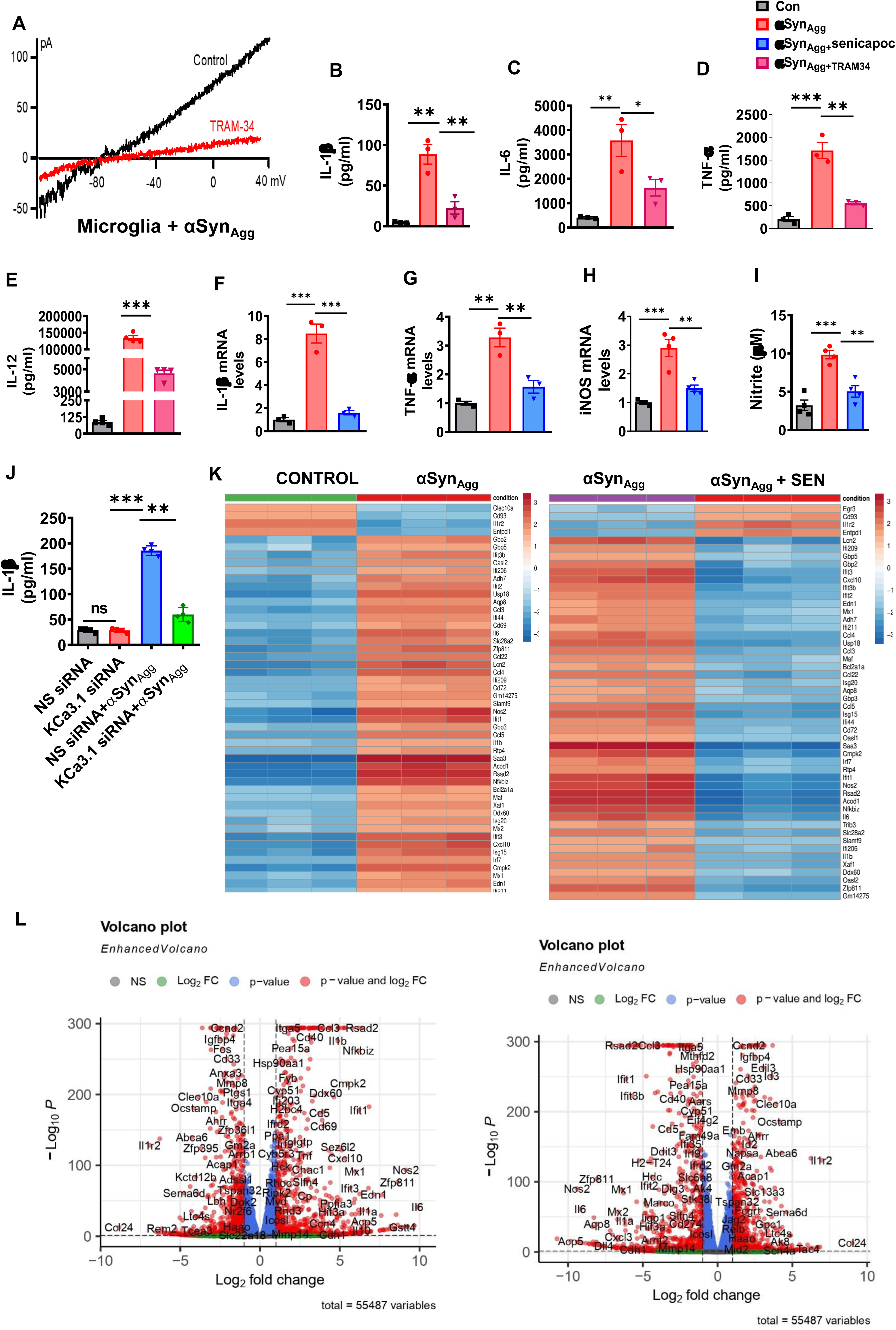
Pharmacological inhibition of KCa3.1 suppresses channel activation and alters the whole-cell proteomics profile in αSyn_Agg_-treated microglial cells. (A) Patch-clamp analysis of KCa3.1 channel activity in αSyn_Agg_-treated primary microglia in the presence or absence of 250 nM TRAM-34. (B-E) LUMINEX ELISA multiplex assay of pro-inflammatory cytokine release in primary microglia pre-treated with 1 μM TRAM-34 for 1 h followed by exposure to 1 μM αSyn_Agg_ for another 24 h. Secreted cytokines determined in the culture supernatants included (B) IL-1β, (C) IL-6, (D) TNF-α, and (E) IL-12. (F-H) RT-qPCR analysis of primary cultured microglia treated with 1 μM αSyn_Agg_ in the presence or absence of the KCa3.1-specific inhibitor senicapoc (5 µM) for 24 h showing inhibited expression of (F) IL-1β, (G) TNF-α, and (H) iNOS. (I) Nitrite quantification by Griess assay. (J) ELISA analysis of IL-1β levels in the culture supernatants of primary microglial cells cultured for 3 d after siRNA-mediated knockdown of KCa3.1 followed by treatment with αSyn_Agg_ for another 24 h. (K) A heat map generated by hierarchical clustering using the relative expression of each probe set is indicated by color intensity, where green indicates lower expression and red indicates higher expression in control, αSyn_Agg_, and αSyn_Agg_ + senicapoc. (L) Volcano map shows the top 50 differentially expressed genes in control, αSyn_Agg_, and αSyn_Agg_ + senicapoc, the abscissa is log2 fold change and the ordinate is -log10 (p-value). The two vertical dotted lines in the figure are the two-fold difference thresholds; the horizontal dotted line is the threshold, p-value = 0.05. (red, increase; blue, decrease). Data are mean ± SEM with 3-6 biological replicates from 3-4 independent experiments unless otherwise noted. Student’s t-tests or one-way ANOVA with Bonferroni multiple comparison tests were utilized for statistical analysis. *p≤0.05, **p<0.01, *** p<0.001.

### Aggregated αSyn treatment provokes a pro-inflammatory phenotype via a Kca3.1-dependent mechanism in primary microglial cells in culture

To better understand what protein pathways KCa3.1 channels may regulate under microglial inflammation, a mouse microglial cell line (MMC) [36–39] was exposed to αSyn_Agg_ and the KCa3.1 inhibitor, TRAM-34, for 24 h and subsequently whole-cell proteomics was carried out on the cell lysates. We identified 169 proteins that were significantly altered between αSyn_Agg_-exposed microglia and controls. When comparing those 169 proteins differentially altered by αSyn_Agg_, 24 of the proteins that were modulated by αSyn_Agg_ were modified by TRAM-34 co-treatment (Fig. S1E-G). Based on this comparison, we analyzed the 24 mutually regulated proteins with STRING, and found 3 significantly altered biological processes, including ncRNA processing, rRNA processing, and TLR4 pathway regulation (Fig. S1H). Both ACOD1 and CD14 are implicated in TLR4 signaling, with CD14 functioning as a positive regulator and ACOD1 as a negative regulator. Furthermore, TLR4 signaling has been implicated in αSyn-induced activation of microglia and astrocytes [40, 41]. This suggests KCa3.1 functions as a sensitive rheostat of canonical or classical pro-inflammatory signaling in microglia after α-synucleinopathy.

To further evaluate the impact of αSyn_Agg_ on the microglial activation response, we carried out transcriptomic profiling of primary microglial cells that were treated with either monomer (con) or αSyn_Agg_ in the presence or absence of the KCa3.1 inhibitor senicapoc at 12 h post stimulation. Initially, we examined the differentially expressed genes (DEGs) between control and αSyn_Agg_ with respect to senicapoc. We examined the top 50 upregulated genes in our RNA-seq datasets in αSyn_Agg_-treated cells as compared to monomer and αSyn_Agg_/senicapoc-treated cells. Upon analysis of the DEGs using GO enrichment analysis, our results reveal that several of the upregulated genes were associated with the regulation of the TLR4 pathway and pro-inflammatory innate immune markers. These overlapping upregulated DEGs, including ACOD1, CCL3, CCL24, CCL4, CCL5 and LCN2 are primarily associated with inflammatory responses [42–44]. In line with our whole cell proteomics analysis (Fig. S1D-H), increased ACOD1 expression was positively correlated with the other pro-inflammatory neurotoxic mediators, such as IL-6, IL1-β and NOS2, suggesting that it is most likely to play a pro-inflammatory role in αSyn_Agg_-induced reactive microgliosis. Additionally, gene expression of IFN-regulated genes, such as Ifi44, Ifi209, Ifit1, Ifi206 and Ifit3b, was significantly higher in microglia exposed to αSyn_Agg_, as compared to monomer-treated cells. Furthermore, inflammatory transcription factors such as Ifn7, Nfkbiz, Saa3, Bcl2a1a and Rsad2, which are positive regulators of neuroinflammation, were strongly increased in the gene expression profile (Fig. 2K-L). Importantly, we found a parallel increase in Cmpk, which is the positive regulator of NLRP3 inflammasome activation [45]. Conversely, senicapoc treatment ameliorated αSyn_Agg_-induced upregulation of proinflammatory mediators in PMG. Remarkably, upregulation of ACOD1, a mitochondrial enzyme that catalyzes itaconate, was evidenced in primary microglia stimulated with αSyn_Agg_, consistent with a previous study from our group [37]. In accordance with our finding, proinflammatory MYD88/STING1-dependent ACOD1 activation was implicated in sepsis-induced injury [41]. Additionally, the cGAS sting pathway has been linked to neuroinflammation-associated neurodegeneration in an α-synucleinopathy mouse model of PD [46], further supporting the pivotal role of proinflammatory mediators in aggravating PD-like neuropathology. As shown above, a partial list of aforementioned proinflammatory genes that are regulated by KCa3.1 was further validated by qPCR analysis (Fig. 2F-H). Collectively, these findings highlight the detrimental role of KCa3.1 in promoting a proinflammatory microglial phenotype in response to αSyn_Agg_ and that the oxidative stress-mediated NLRP3/IL-1β-driven innate immune pathway may in part contribute to neuroinflammation-associated nigral DAergic neurodegeneration in the context of α-synucleinopathy.

### Aggregated αSyn promotes KCa3.1 upregulation via Fyn-mediated STAT1 activation

To determine whether STAT1 regulates *KCNN4* (KCa3.1 gene) expression through binding to its promoter, we determined transcription factor binding sites for STAT1 in KCNN4 regulatory regions using the JASPAR database. As expected, multiple binding sites for STAT1, including the first 500-bp promoter region upstream of the transcription start site on the KCCN4 gene promoter, were discovered, suggesting that STAT1 can regulate KCNN4 expression (Fig. 3A). Indeed, using chromatin immunoprecipitation (ChIP) analysis, we demonstrated that αSyn_Agg_ treatment significantly increased the recruitment of STAT1 to the KCNN4 promoter as compared to controls, indicating that KCNN4 is transcriptionally regulated by STAT1 (Fig. 3B) [47]. Next, we explored whether αSyn_Agg_ treatment induces STAT1 activation. Our WB analysis revealed an increase in STAT1 expression in primary microglial cells treated with αSyn_Agg_ (Fig. 3C). Additionally, our immunofluorescence studies revealed a prominent nuclear translocation of STAT1 (Fig. 3D). To further verify that STAT1 regulates KCa3.1 expression, we used a cell-permeable inhibitor of STAT1, fludarabine. As anticipated, we observed a reduction in the mRNA expression of the STAT1-targeted genes TNF-α (Fig. 3E) and IL-1β (Fig. 3F), as well as the protein (Fig. 3G) and mRNA (Fig. 3H) expression of KCa3.1 in αSyn_Agg_-stimulated mouse microglial cells (MMCs), indicating the stimulatory effects of STAT1 on KCa3.1 activation and associated STAT1-targeted genes in response to αSyn_Agg_ [38]. In addition, MMCs transfected with STAT1 siRNA showed reduced KCa3.1 expression (Fig. 3I) and reduced mRNA expression of the proinflammatory cytokines IL-1β and TNF-α (Fig. 3J). Given that we have previously demonstrated that Fyn activation in reactive microglia is linked to neuroinflammation and associated nigral neurodegeneration in experimental models of PD through a transcription factor-dependent manner [21, 48], we next investigated whether primary microglia harvested from conventional Fyn KO mice exhibit reduced STAT1 activation in response to αSyn_Agg_. Our WB analysis revealed reduced STAT1 phosphorylation in Fyn KO microglia as compared to WT primary microglia stimulated with αSyn_Agg_ (Fig. S2). These findings demonstrate that a functional interplay between Fyn and STAT1 in reactive microglia promotes KCa3.1-mediated reactive microgliosis in response to αSyn_Agg_.

**Fig 3.**
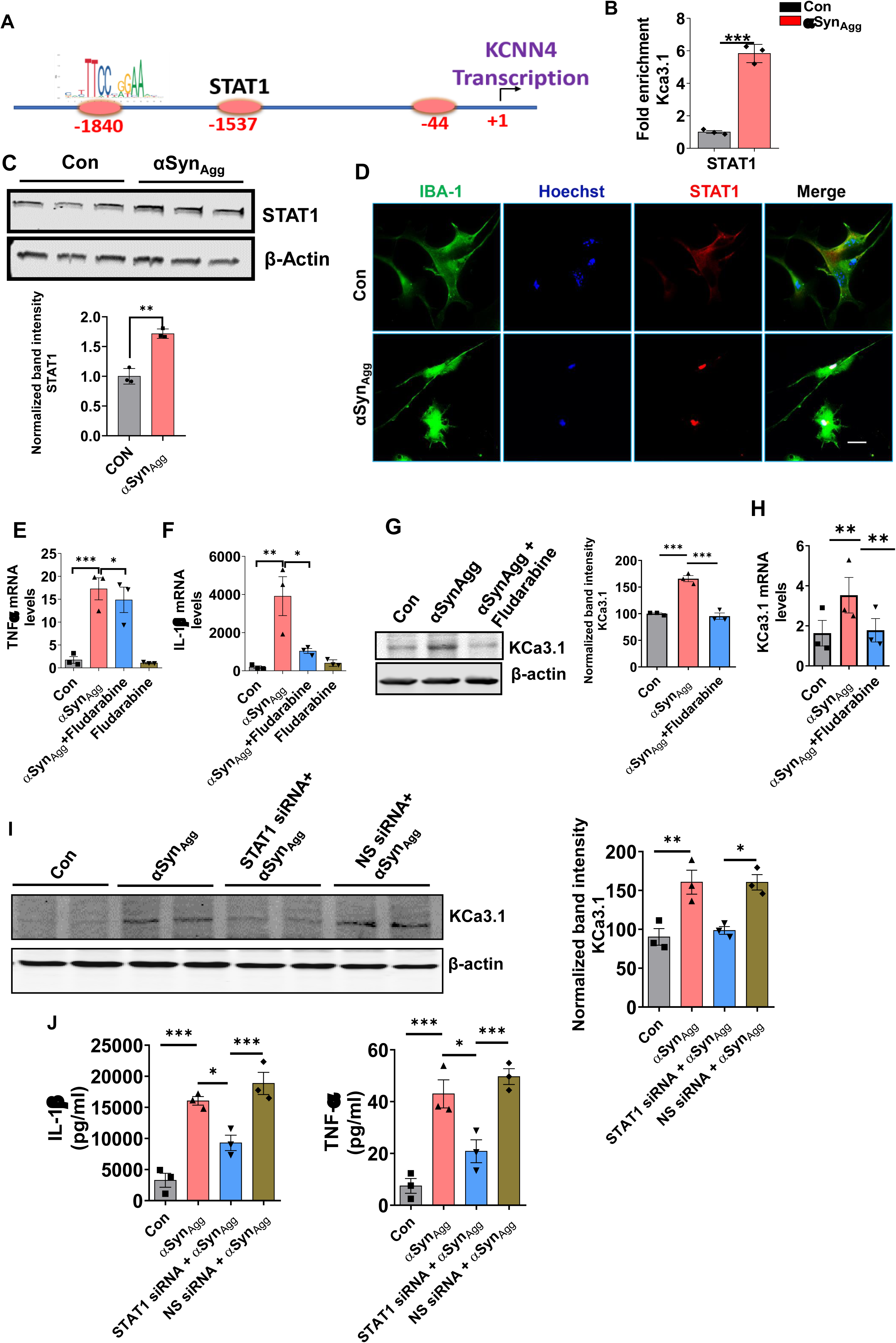
STAT1 regulates KCa3.1 expression. (A) Schematic representation of mouse KCa3.1 gene promoter. (B) ChIP analysis of GAS/ISRE element within the KCa3.1 promoter region showing Increased binding of STAT1 to the proximal promoter region of KCa3.1 in αSyn_Agg_-treated primary microglia. Quantification of PCR product is shown. Data is representative of at least 3 independent experiments. (C) Western blot (WB) analysis of STAT1 expression in primary microglial cultures treated with or without αSyn_Agg_ for 24 h. (D) Representative immunofluorescence images showing increased nuclear localization of STAT1 in primary microglia stimulated with 1 μM αSyn_Agg_ for 3 h (n=3). Scale bar, 15 µm. (E-H) Mouse microglial cells (MMCs) pre-treated with fludarabine for 1 h, followed by 1 μM αSyn_Agg_ for another 24 h were assayed for (E-F) TNFα and IL-1β mRNA expression using RT-qPCR analysis, (G) KCa3.1 protein expression using WB, and (H) KCa3.1 mRNA expression. (I) Immunoblotting KCa3.1 in primary microglial cell lysates transfected with either scrambled siRNA (NS siRNA) or STAT1 siRNA, and exposed to αSyn monomer or αSyn_Agg_ (1 µM) for another 24 h after the 72-h transfection phase. (J) RT-qPCR analysis showing fold change in mRNA expression of STAT1-regulated inflammatory genes in primary microglia cells with or without STAT1 siRNA knockdown in the presence or absence of αSyn_Agg_ (n=2). Data are expressed as the means ± SEM; n=3 per group. *p≤0.05, **p<0.01 and ***p<0.001.

### Fyn kinase promotes KCa3.1 expression and the proinflammatory activation phenotype of microglia in diverse models of PD

We previously reported increased expression of Fyn in the SNpc of PD patients as well as in mouse models of α-synucleinopathy; however the relationship between FYN activation and a-synucleinopathy remains poorly characterized [21]. The above experiments demonstrated a role for STAT1 in the regulation of KCa3.1 expression in a Fyn kinase dependent manner in response to αSyn_Agg_. In the current study, we aimed to determine if FYN kinase induces changes in KCa3.1 expression and associated downstream signaling mediators. We next evaluated its effects on KCa3.1 upregulation in response to Syn PFF. We confirmed that stimulation of primary murine microglia with either LPS or αSyn_Agg_ induced a time-dependent increase in the activation of Fyn as reflected by increased Fyn phosphorylation (Fig. 4A). Moreover, to validate the expression of activated Fyn in adult microglial cells stimulated with αSyn_Agg_, we immunostained for IBA-1 and pFyn. Consistent with our previous study, we found a dramatic increase in the area covered by pFyn within IBA-1-positive cells, which was accompanied by marked changes in the microglial morphology including reduced process length and increased area of the soma (Fig. 4B). Additionally, using in situ proximity ligation assay (PLA), we investigated whether Fyn might bind to KCa3.1. We detected more than a 4-fold increase in Fyn/KCa3.1 interactions (Fig. 4C), as indicated by PLA dots in αSyn_Agg_-treated cells as compared to control cells, supporting the physical interaction between Fyn and KCa3.1 in the context of α-synucleinopathy.

**Fig 4.**
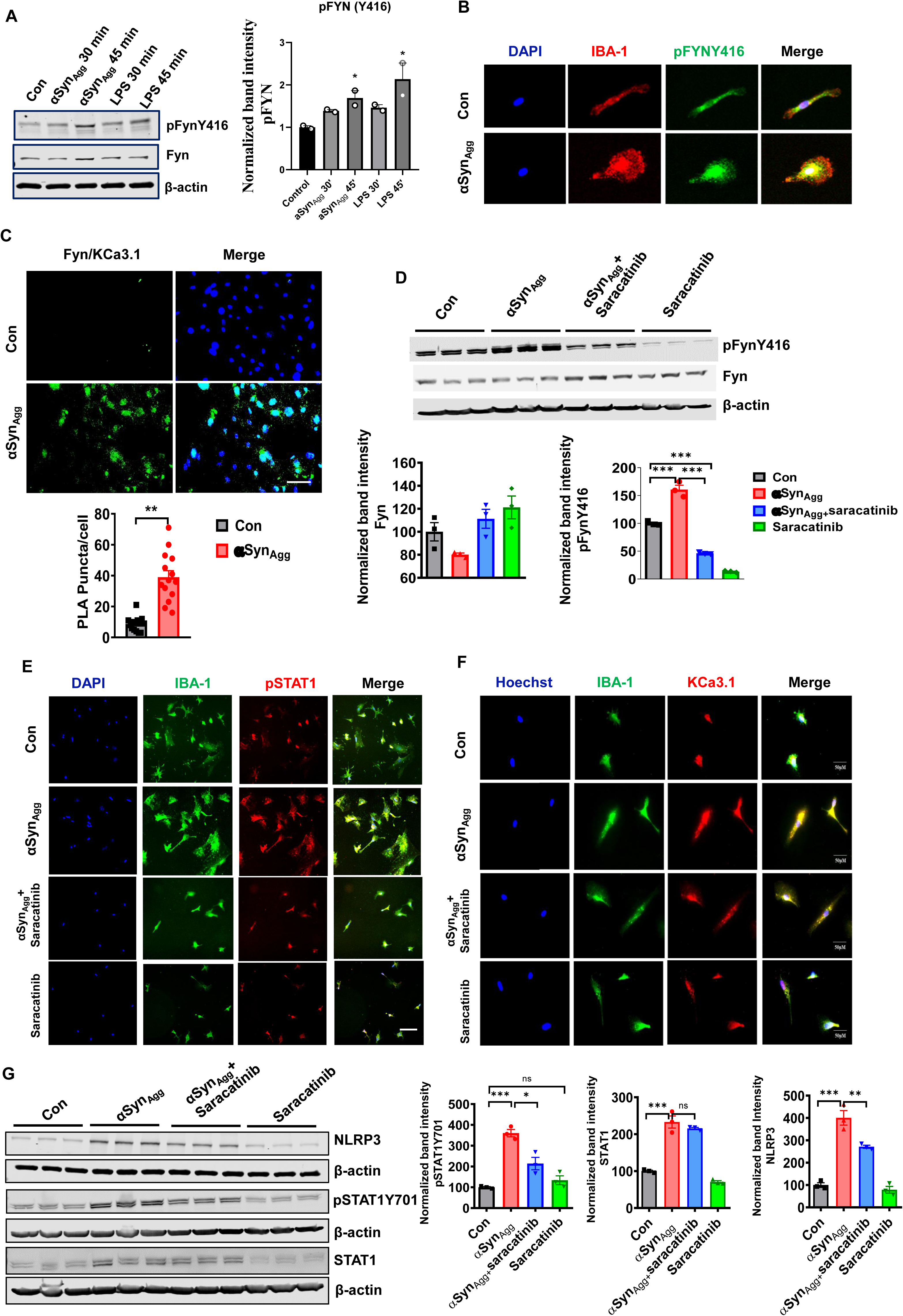
Pharmacological blockade or genetic ablation of Fyn suppresses reactive microglia phenotype in MMC and primary microglial cells treated with either LPS or αSyn_Agg_. (A) Immunoblot analysis of both pFynY416 and native Fyn activation in adult WT mouse primary microglia treated with 1 μM αSyn_Agg_ or LPS (100 ng/mL). Right: Quantification of pFynY416 signal intensity normalized to native Fyn. (B) Representative double immunofluorescence images displaying IBA-1 (red) and pFynY416 (green) showing membrane translocation accompanied by marked nuclear localization of pFynY416. Nuclei were counterstained with DAPI (blue). Scale bar, 50 µm. (C) Proximity ligation assay (PLA) of interaction between Fyn and KCa3.1 in αSyn_Agg_-stimulated primary microglia showing Fyn-KCa3.1 proximity ligation signal (green) and DAPI (blue). Representative immunofluorescence images from 3-4 independent samples. The relative intensity of Fyn-KCa3.1 PLA signals (per DAPI signal) was quantified and is displayed in the bottom panel. Scale bar, 10 µM. (D) Representative Western blot (WB) images showing reduction in pFynY416 expression in αSyn_Agg_-stimulated MMC cells pre-treated with saracatinib (5 µM). Bottom: densitometric quantification of pFynY416 and native Fyn (n=3). (E-G) Primary microglial cells treated with αSyn_Agg_ (1 µM) in the presence or absence of saracatinib (5 µM) for 24 h. (E) Representative double immunolabeling images for STAT1 and IBA-1. (F) Representative immunofluorescence image showing reduced KCa3.1 expression in the presence of both αSyn_Agg_ and saracatinib for 24 h. Double immunolabeling fluorescence analysis was performed using antibodies against IBA-1 (green) and KCa3.1 (red). Scale bar = 50 µm. (G) WB analysis of primary microglial lysates that were probed with antibodies against STAT1, pSTAT1Y701 and NLRP3. Data are presented as mean ± SEM. n=3-4 per group. *p≤0.05, **p<0.01, ***p<0.001, determined by One-way ANOVA followed by Dunnett’s multiple comparison test. ns, not significant.

Next, using primary murine microglial cultures, we investigated whether pharmacological inhibition of Fyn via saracatinib (AZD0530), a centrally active Fyn/Src inhibitor, ameliorates Fyn kinase activation and associated inflammatory mediator generation. In agreement with a direct functional role of FYN kinase activation in reactive microgliosis our immunoblotting studies revealed increased expression of activated Fyn kinase in αSyn_Agg_-only-treated cells, and this increase was markedly reduced by saracatinib pre-treatment prior to αSyn_Agg_ stimulation, indicating target (FYN kinase) engagement by Saracatinib(Fig. 4D). Furthermore, we found that saracatinib also reduced αSyn_Agg_-induced upregulation of NLRP3, KCa3.1, and STAT1 protein expression (Fig. 4E-G), concomitant with proinflammatory cytokine generation such as IL-1β, TNF-α and IL-6 (Fig. S3A), thereby supporting a role for Fyn in the regulation of Kca3.1 via a ATAT1/NLRP3 dependent signaling axis in α-synucleinopathy.

To further analyze the contribution of Fyn kinase to KCa3.1-mediated reactive microgliosis, we investigated the inflammatory response of mouse primary microglia from WT and Fyn KO mice treated with αSyn_Agg_. Our results indicate that, compared to Fyn^+/+^ primary microglia, Fyn KO primary microglial cultures exhibited significantly reduced KCa3.1 mRNA and protein upregulation after stimulation with αSyn_Agg_ (Fig. 5A-C), suggesting that Fyn is involved in KCa3.1 upregulation in response to αSyn_Agg_. Next, we investigated the role of Fyn in *in vivo* models of neuroinflammation. As anticipated, intracerebroventricular (ICV) injection of LPS significantly increased the mRNA expression of proinflammatory mediators, including IL-1β, TNF-α, IL-6 and iNOS (Fig. 5D-G) in the striatum. To provide additional support for the role of Fyn in reactive microgliosis we generated Fyn(fl/fl): Cx3cr1-GFP-Cre (Tg/0) mice that exhibit genetic depletion of Fyn specifically in the microglia (Fig. S3B-C). As expected, the mRNA expression of IL-1β, TNF-α, and Nos2 was reduced in isolated adult microglia of LPS-treated Fyn cKO mice as compared to littermate controls (Fig. S3D-F). Next, we assessed the reactive microglial phenotype by determining the extent of amoeboid reactive microglia in LPS-treated Fyn cKO mice. Our IHC studies reveal that treatment of Fyn cKO mice with LPS led to an amelioration of hypertrophic microglia with the loss of ramified branches as compared to littermate controls (Fig. S3G), suggesting a proinflammatory role for microglia Fyn in response to LPS. Consistent with a proinflammatory role for microglial fyn in reactive microgliosis. Additionally, a marked increase in the expression of cortical IL-1b, TNF alpha, iNOS alongside NLRP3 was evidenced as assessed via qRT PCR analysis while a marked reduction in the expression of afore mentioned mediators was evidenced in LPS treated Fyn cKO mice Together, these results support a model whereby FYN activation is a critical contributor to reactive microgliosis, which in turn converges upon KCa3.1 in a STAT1 dependent manner to augment reactive microglial phenotype. Together, these data demonstrate that pharmacological and microglia-specific deletion of FYN might be responsible for KCa3.1 mediated reactive microgliosis via a STAT1/NLRP3 dependent signaling axis.

**Fig. 5.**
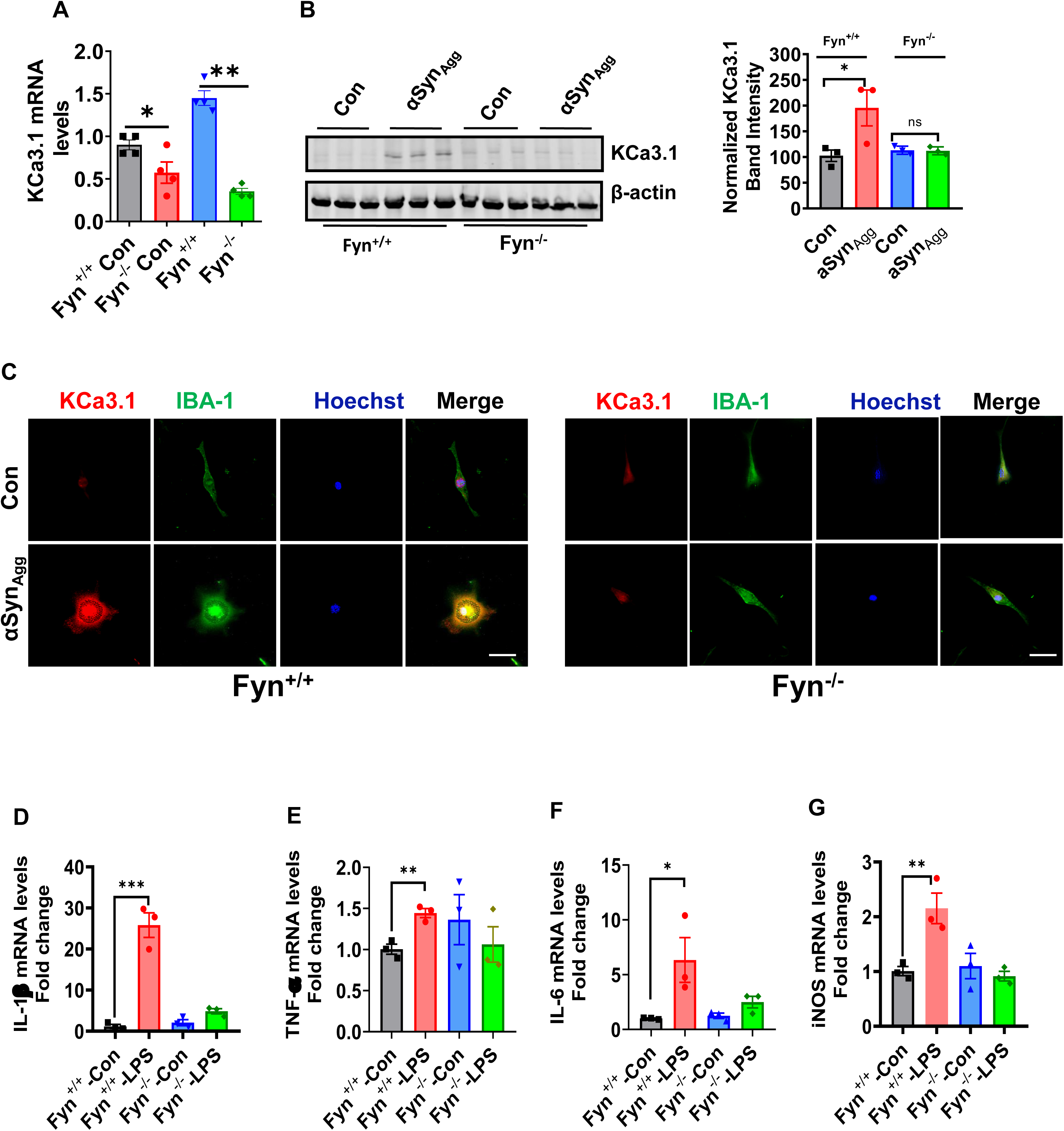
Fyn deficiency ameliorates LPS-induced KCa3.1 activation and proinflammatory cytokines upregulation in cultured microglial cells. Fyn deficiency attenuates αSyn_Agg_-induced KCa3.1 (A) mRNA and (B) protein expression in primary microglial cultures. (C) Representative immunofluorescence images showing reduced KCa3.1 immunoreactivity in Fyn^-/-^ primary microglia. (D-G) Expression of (E) IL-1β, (F) TNF-α, (G) IL-6, and (H) iNOS from WT and Fyn^-/-^ primary microglia treated with vehicle or LPS for 24 h. Data are presented as mean ± SEM (n=3-4 per group). One-way ANOVA followed by Tukey’s post hoc test. *p≤0.05, **p<0.01, ***p<0.001.

### Pharmacological blockade of KCa3.1 via senicapoc reduces nigral DAergic neurodegeneration and immune dysregulation in MPTP-treated mice

Having demonstrated the therapeutic efficacy of senicapoc in an AD mouse model, and that it exhibits excellent BBB permeability [30], next we examined the effect of senicapoc on M1 and M2 markers in the brains of MPTP-treated mice. Proinflammatory cytokines such as TNF-α and IL-1β have been linked to nigral DAergic neurodegeneration in PD as well as multiple models of neurodegeneration [49]. The nigral brain region of senicapoc-treated MPTP mice displayed decreased levels of TNF-α and IL-1β as compared with MPTP alone mice (Fig. 6A-B). In contrast, senicapoc ameliorated MPTP-induced TGFβ and IL-10 depletion, though the effect on IL-10 was not statistically significant, while no changes in ARG-1 and CD68 protein expression were observed (Fig. 6C). Our findings are in accordance with previous studies demonstrating the anti-inflammatory effects of senicapoc in neurodegenerative models [30, 50]. We next evaluated the signaling pathways potentially linked to senicapoc-mediated DAergic neuroprotection. As anticipated, senicapoc treatment diminished MPTP-induced STAT1 activation as well as KCa3.1 upregulation, and restored TH expression, suggesting that the KCa3.1-associated, STAT1-mediated proinflammatory signaling cascade may contribute to MPTP-induced DAergic neurotoxicity (Fig. 6D) [51]. Next, we examined to what extent reduced neuroinflammation in MPTP/senicapoc mice impacted nigral DAergic neurodegeneration. At 11 d post-MPTP administration, unbiased stereological analysis revealed a significant loss of TH-positive nigral neurons in the substantia nigra of MPTP-treated mice, while senicapoc treatment afforded partial protection against MPTP-induced nigral DAergic neurodegeneration (Fig. 6E). Furthermore, the partial rescue of nigral DAergic neurons by senicapoc in MPTP-treated mice closely paralleled partial restoration of DA and its metabolite 3,4-dihydroxyphenyacetic acid (DOPAC) levels (Fig. 6F-G). To further determine whether intraperitoneal administration of senicapoc was able to attain therapeutic concentrations within the MP mouse brain, we treated a subset of MP mice with LPS (3 mg/kg) [52, 53] followed by senicapoc treatment at 40 mg/kg [30] for another 2 wk, and at the end of the treatment period, cortical brain regions were collected for pharmacokinetic analysis. Ultra high-pressure liquid chromatography (UPLC)/mass spectrometry (MS) analysis of cortical brain samples demonstrated that total senicapoc brain concentrations varied between 4 and 9 µM, which is 8 times above the IC_50_ value of 11 nM, suggesting that the brain concentration was sufficient to elicit a pharmacodynamic effect [30, 50] (Fig. S4A). The increase in brain concentration of senicapoc positively correlated with a reduction in LPS-induced NLRP3 levels, although the trend was not statistically significant, highlighting its anti-inflammatory effects in PD mouse models (Fig. S4B). Indeed, a previous publication demonstrated that NLRP3 KO mice exhibit resistance against MPTP-induced neuroinflammation and associated nigral DAergic neurodegeneration [54], further corroborating the pivotal role of the NLRP3-mediated innate immune response in PD-associated nigral DAergic neurodegeneration. Thus, limiting KCa3.1 activation via senicapoc may represent a novel strategy to suppress neuroinflammation caused by PD-related neurotoxins.

**Fig 6.**
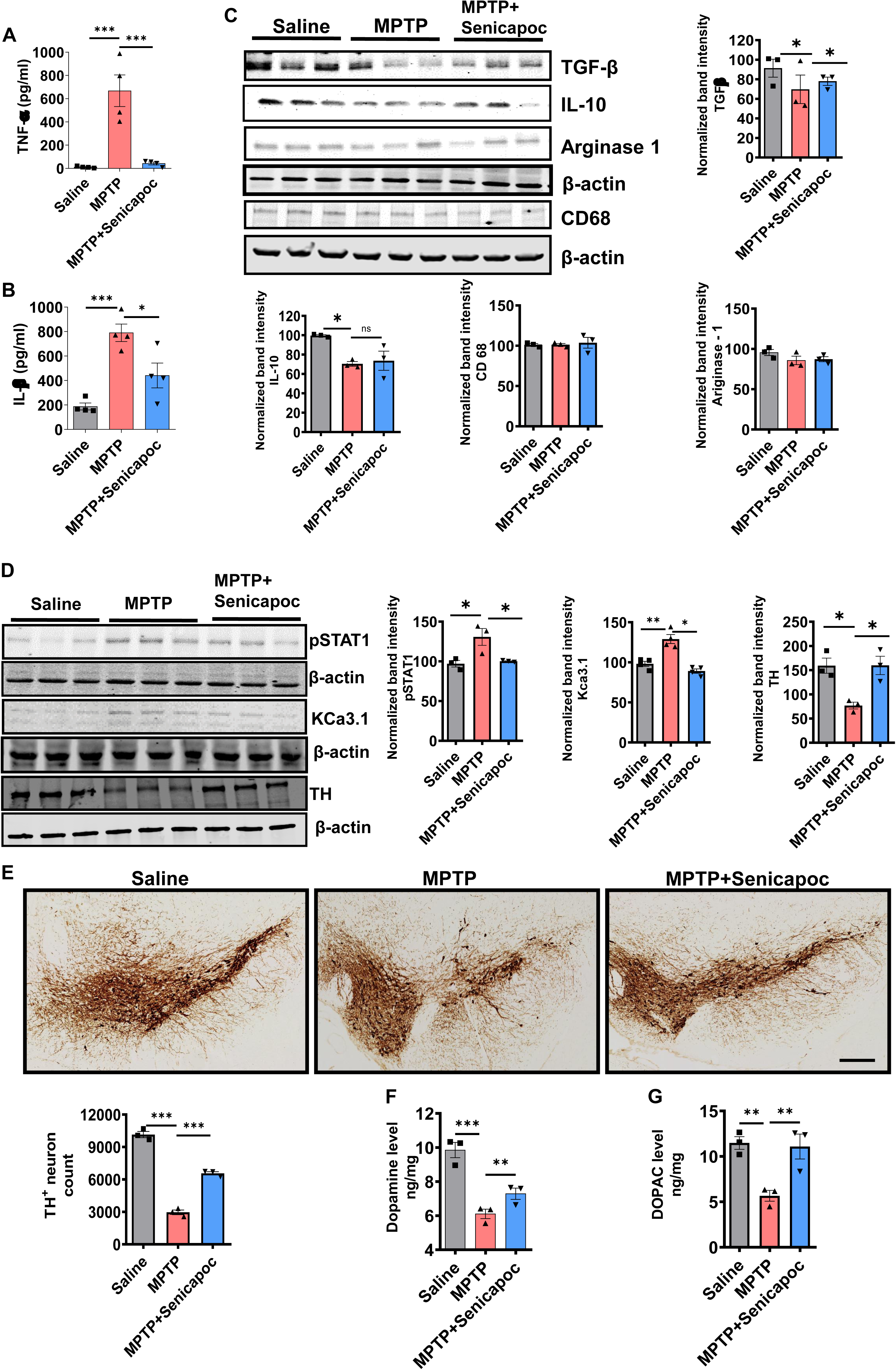
Pharmacological blockade of KCa3.1 via senicapoc attenuates PD-like pathology in an MPTP mouse model of PD. LUMINEX multiplex analysis of the levels of the proinflammatory cytokines (A) TNF-α and (B) IL-1β in the substantia nigra following MPTP treatment in the presence or absence of senicapoc (n=4 mice/group). (C) Representative Western blots (WBs) for TGF-β, IL-10, Arginase 1 and CD68 and their densitometric quantification normalized to β-actin (lower panels) depicting protein expression 7 d post-MPTP in the SNpc of MPTP-treated mice that either received vehicle or senicapoc. (D) Immunoblot analysis showing levels of pSTAT1, KCa3.1, and TH and (right panels) their quantification normalized to β-actin. (E) Representative TH immunohistochemical imaging studies of the midbrain coronal sections of MPTP- and saline-treated mice receiving senicapoc or vehicle. Scale bar: 200 µm. Bottom left: Stereological assessment of TH-positive neurons in midbrain coronal sections (n = 3-4). (F-G) HPLC analysis of striatal DA and DOPAC levels in saline- and MPTP-treated mice that either received senicapoc or vehicle (n=3). Data represented as mean ± SEM from 3 mouse brains. Significance was determined by one-way ANOVA followed by a Tukey’s post hoc test. *p≤0.05, **p<0.01, ***p<0.001.

### Pharmacological blockade of KCa3.1 via TRAM-34 reduces nigral DAergic neurodegeneration and neuroinflammation in an α-synucleinopathy mouse model

To further verify the role of KCa3.1 in regulating αSyn_Agg_-induced neuroinflammation and associated nigral DAergic neurodegeneration, we evaluated the therapeutic efficacy of another KCa3.1 inhibitor, TRAM-34, in a chronic progressive model of α-synucleinopathy. For this purpose, we used the intrastriatal αSyn preformed fibril (PFF) model of PD, whereby a single injection of αSyn_PFF_ instilling recombinant αSyn proteins into various brain regions in different rodent models has been shown to result in widespread αSyn pathology in the interconnected brain regions [55–57]. Previous studies have demonstrated that αSyn_PFF_-inoculated mice progressively exhibit nigral DAergic neuronal degeneration and associated behavioral deficits [58]. Moreover, motor dysfunction originating from nigrostriatal DAergic neurodegeneration and DA depletion has been identified as key hallmarks of PD [59]. After 28 wk following intrastriatal αSyn_PFF_ inoculation, mice from different treatment groups including αSyn_PFF_- and αSyn_PFF_/TRAM-34-treated mice were tested for behavioral deficits using the Versamax locomotor activity monitor. Representative locomotor activity maps of movement of αSyn_PFF_-treated mice in the presence or absence of TRAM-34 are shown in Fig. 7A. A significant improvement in motor functions was observed in αSyn_PFF_/TRAM-34-treated mice compared to αSyn_PFF_-treated mice, as quantified by improved behavioral parameters including horizontal activity (Fig. 7B), total distance travelled (Fig. 7C), total rest time (Fig. 7D) and ambulatory activity count (Fig. 7E), suggesting that inhibition of KCa3.1 can rescue locomotor deficits in the α-synculeinopathy mouse model of PD. Previous studies have reported a progressive loss of DAergic neurons in the substantia nigra concomitant with αSyn pathology on the injected side [60]. In line with these studies, we found a significant reduction in striatal DA and DOPAC levels at 6 mo post-αSyn_PFF_ injection. In contrast, TRAM-34-treated mice exhibited a remarkable recovery of DA and DOPAC levels (Fig. 7F-G), indicating that blockade of KCa3.1 may serve as a novel therapeutic avenue in PD treatment. We next investigated the impact of TRAM-34 on αSyn_PFF_-induced TH neuronal loss. WB further revealed that TRAM-34-treated αSyn_PFF_ mice exhibited a nonsignificant trend toward increased TH expression as compared to αSyn_PFF_-treated mice (Fig. 7H). Additionally, our IHC studies further confirmed the preservation of nigral DAergic neuronal integrity in TRAM-34-treated αSyn_PFF_ mice as compared with αSyn_PFF_-treated mice (Fig. 7I). To further confirm if the DAergic neuroprotection observed in TRAM-34-treated αSyn_PFF_ mice is attributed to reduced inflammatory response, we assessed the levels of proinflammatory cytokines in the serum at 5 mo post-αSyn_PFF_ inoculation.Next we assessed the effects of αSyn_PFF_ inoculation on serum proinflammatory cytokines. αSyn_PFF_ induced an upregulation of serum proinflammatory cytokines including IL-6 and IL-1β as compared to saline-treated controls (Fig. 7J). In contrast, TRAM-34 treatment ameliorated αSyn_PFF_-induced upregulation of proinflammatory cytokines, suggesting that blockade of the αSyn_PFF_-induced peripheral proinflammatory response through KCa3.1 inhibition may in part contribute to DAergic neuroprotection in TRAM-34-treated αSyn_PFF_ mice. These data demonstrate that TRAM-34 may also serve as a therapeutic target in PD and related α-synucleinopathies.

**Fig 7:**
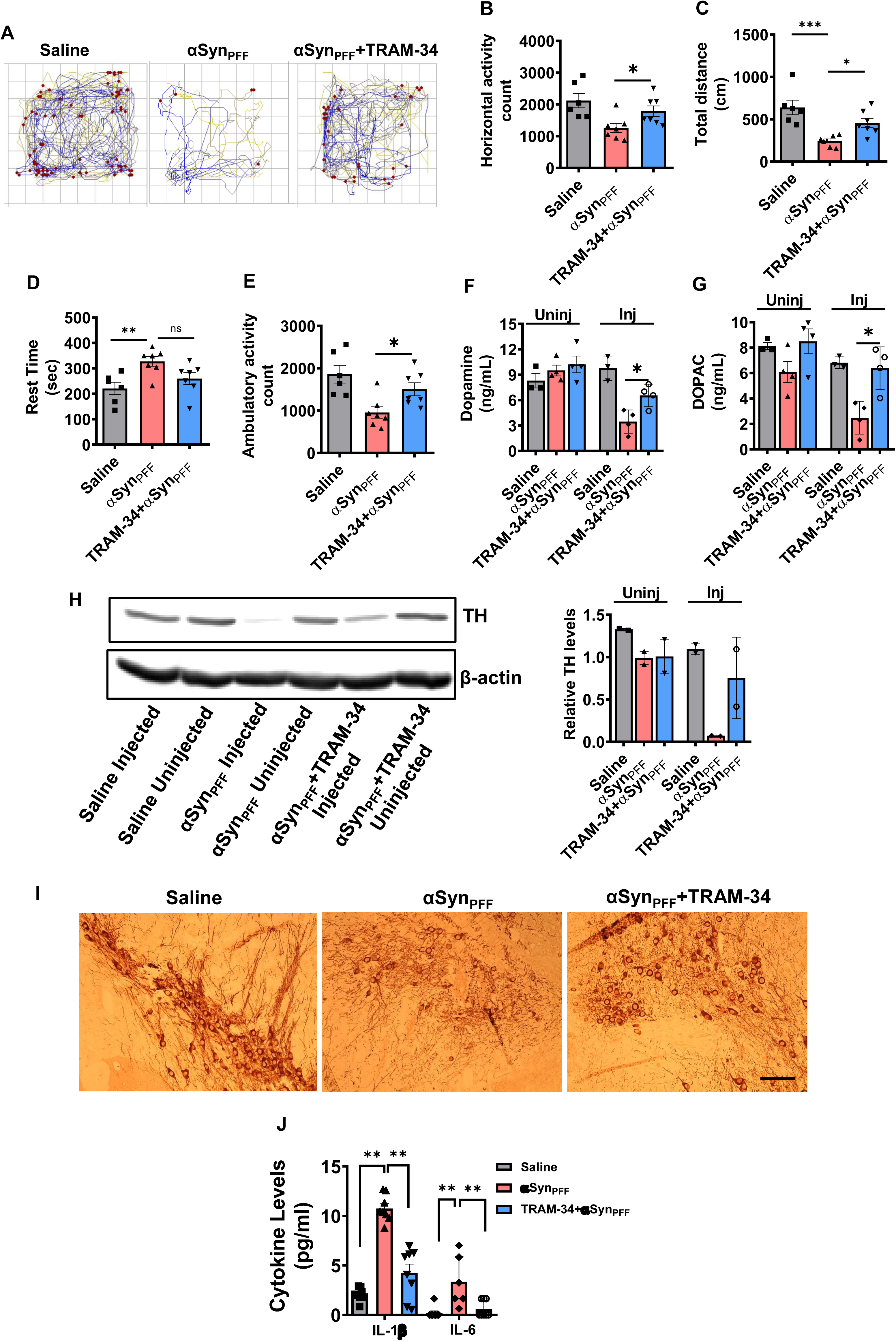
Pharmacological blockade of KCa3.1 via TRAM-34 ameliorates αSyn_PFF_-induced neurobehavioral deficits and loss of nigral TH functionality. (A) Representative movement track of mice. (B-E) Open-field test results showing (B) horizontal activity, (C) total distance travelled, (D) total rest time, and (E) ambulatory activity count. (F-G) HPLC analysis demonstrating the effects of TRAM-34 treatment on (F) DA and (G) DOPAC levels in intrastriatal αSyn_PFF_-inoculated mice. (H) Western blotting analysis and quantification of αSyn_PFF_-induced TH loss in the SNpc brain region that either received saline or TRAM-34. (I) Representative TH immunohistochemical imaging studies on midbrain coronal sections from αSyn_PFF_-treated mice treated with or without TRAM-34. Scale bar: 100 µm. (J) Luminex analysis of plasma cytokine levels of IL-1β and IL-6. Data represented as mean ± SEM from 4-6 per group. Significance was determined by one-way ANOVA followed by a Tukey’s post hoc test. *p≤0.05, **p<0.01.

### KCa3.1 deficiency alleviates neuroinflammation and PD-like neuropathology in diverse mouse models of PD

It has previously been reported that inhibition of KCa3.1 by senicapoc confers neuroprotection and diminished reactive microgliosis in AD and LPS-treated mouse model [30]. We reasoned that if KCa3.1 is a central mediator of αSyn_Agg_-mediated neuroinflammation and associated nigral DAergic neurodegeneration, then genetic ablation of KCa3.1 could ameliorate hαSyn-AAV-induced Parkinsonism. Morphometric analysis of microglia in the nigral brain sections from αSyn-AAV mice at 2 mo post-AAV inoculation displayed characteristics of a reactive amoeboid microglial phenotype, as assessed via the measurement of branched endpoints, cell body diameter, number of branches, endpoints and average branch length. Remarkably, KCa3.1 KO displayed reduced reactive microglial manifestations as compared to WT mice that received αSyn-AAV inoculation (Fig. 8A-B). Moreover, αSyn-AAV-inoculated mice displayed a reactive microglial phenotype with accompanying increased expression of KCa3.1 in IBA-1+ microglia as compared to control AAV-inoculated mice at 4 mo post-inoculation (Fig. 8A-B). Additionally, WB shows a significant reduction in the IBA-1 expression in the KCa3.1 KO mice inoculated with αSyn-AAV compared to the control AAV-inoculated mice (Fig. S5). This reactive microglia phenotype closely paralleled the increased mRNA expression of proinflammatory markers including TNF-α, IL-1β, and IL-6 in the nigra at 4 mo post-αSyn-AAV injection. Additionally, alternative activation markers in the αSyn-AAV-associated neuroinflammatory response displayed a significant restoration in the expression of IL-4, although the expression of arginase1 remained unchanged in the αSyn-AAV mice (Fig. 8C), in accordance with Theodore et al. [61]. Similarly, the expression of chemokines that promote migration of leuokocytes and lymphocytes such as CCL2, CCL5, and CCR2 was elevated in the striata of αSyn-AAV mice post-inoculation (Fig. 8D), while KCa3.1 KO partially reduced αSyn-AAV-induced expression of proinflammatory cytokines and chemokines, as well as modulated a subset of alternative markers, suggesting a that KCa3.1 may in part mediated CNS influx of peripheral inflammatory cells and inflammatory cytokine generation (Fig. 8C-D). While Fyn has been implicated in driving both neuroinflammation and nigral DAergic neurodegeneration, its role in KCa3.1-mediated neuroinflammation in the context of α-synucleinopathy remains poorly defined. Consistent with the in vivo neuroinflammation studies, we found STAT1 and Fyn kinase activation as well as iNOS and IBA-1 upregulation in the striata at 6 and 16 wk following αSyn-AAV administration. Conversely, KCa3.1 deficiency markedly reduced the induction of aforementioned oxidative stress-responsive inflammation markers induced by αSyn-AAV administration (Fig. 8E), in accordance with Toyama et al.’s [62] studies demonstrating that KCa3.1 inhibition reduces oxidative stress response in an experimental model of atherosclerosis. Collectively, these results suggest that a functional interaction between KCa3.1 and the Fyn/STAT1 signaling axis may in part promote a reactive microglial phenotype in the context of α-synucleinopathy.

**Fig 8.**
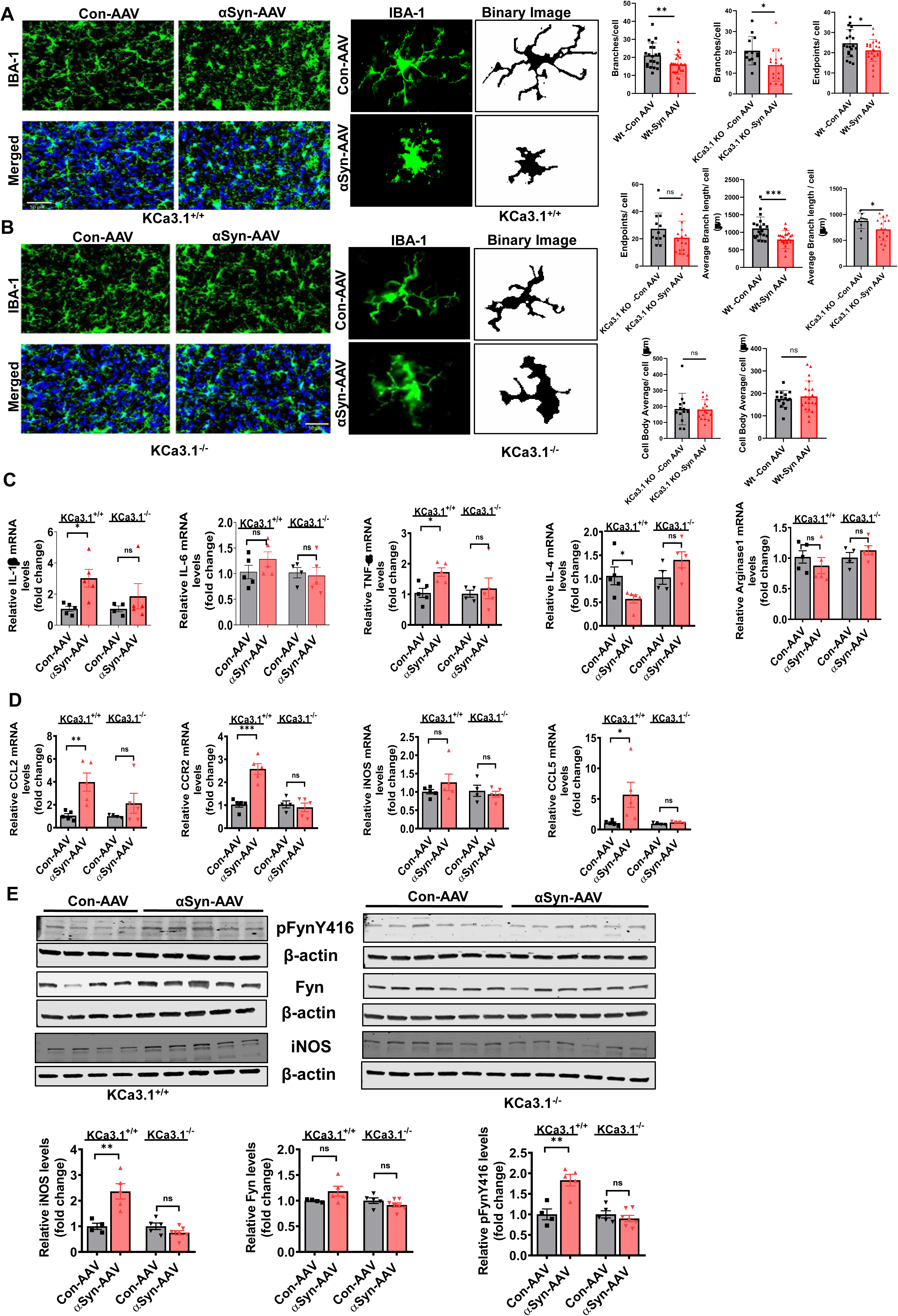
Global knockout of KCa3.1 ameliorates reactive microglial phenotype and PD-like neuropathology in h-αSyn-AAV-inoculated mice. (A) Representative KCa3.1 immunohistochemical images of the striatal coronal sections of KCa3.1 global KO mice and C57 WT controls with or without αSyn-AAV inoculation (n=3-4). Scale bar for high-magnification images (40x): 100 µm. (B) Representative images of individual microglia from the SN region of control and αSyn-AAV-treated groups along with respective skeletonized images. Scale bar: 50 µm. (C-D) RT-qPCR gene expression analysis of proinflammatory mediators showing (C) IL-1β, IL-6, TNF-α, IL-4, Arginase1 and (D) CCL2, CCR2, iNOS, CCL5. (E) Western blot analysis normalized to total Fyn or β-actin showing protein expression of pFynY416, Fyn, and iNOS in the striata of KCa3.1 KO and WT mice with or without αSyn-AAV injection. Significance was determined by one-way ANOVA followed by a Tukey’s post hoc test. *p≤0.05, **p<0.01, ***p<0.001. ns, not significant.

To further test the hypothesis that microglial KCa3.1 is a critical contributor to reactive gliosis, we examined the protein expression of STAT1 and NLRP3, as well as mRNA expression of proinflammatory cytokines such as IL-1β, IL-6, TNF-α in adult microglial cells isolated from KCa3.1 KO and WT mice treated with or without LPS (Fig. S6A) using WB and RT-qPCR analysis. We observed a robust increase in NLRP3 expression concomitant with STAT1 upregulation following stimulation of adult microglial cells with LPS as compared to controls, while KCa3.1 KO adult glial cells exhibited a blunted response to LPS-induced proinflammatory response. Likewise, mRNA expression of IL-1 β, IL-6, and TNF-α was reduced in LPS-treated KCa3.1 KO cells microglia (Fig. S6B). Collectively, these studies suggest that the FYN mediated NLRP3-STAT1 signaling axis converges on KCa3.1 which in turn feeds back on FYN/STAT1 signaling axis further augmenting a-synucleinopathy mediated inflammation.

We further examined the impact of KCa3.1 KO on αSyn pathology. αSyn pathology was examined via WBting analysis using an antibody against hαSyn (Anti-Alpha-synuclein antibody [MJFR1]). As anticipated, administration of αSyn-AAV promoted h-αSyn overexpression in the striata at 6 wk (Fig. 9A). Interestingly, in KCa3.1 KO mice that received αSyn-AAV injection, we observed a marked reduction in the expression αSyn_Agg_ in the striata as compared to WT mice (Fig. 9A). To further validate the formation of pathological αSyn-positive inclusions, we conducted IHC studies using an antibody against phosphorylated αSyn in the ventral midbrain region. We found robust accumulation of αSyn-positive inclusions within the nigral brain regions of αSyn-AAV-inoculated mice (Fig. 9B), in line with Arawaka et al.’s [63] findings. Furthermore, the αSyn-positive inclusions displayed LB-like characteristics including resistance to proteinase K digestion (Fig. 9B) [33]. Interestingly, the expression αSyn-positive inclusions were partially reduced in KCa3.1 KO mice that received the αSyn-AAV injection (Fig. 9B), which suggests a protective effect of KCa3.1 gene deficiency on αSyn pathology in α-synucleinopathy mice.

**Fig 9:**
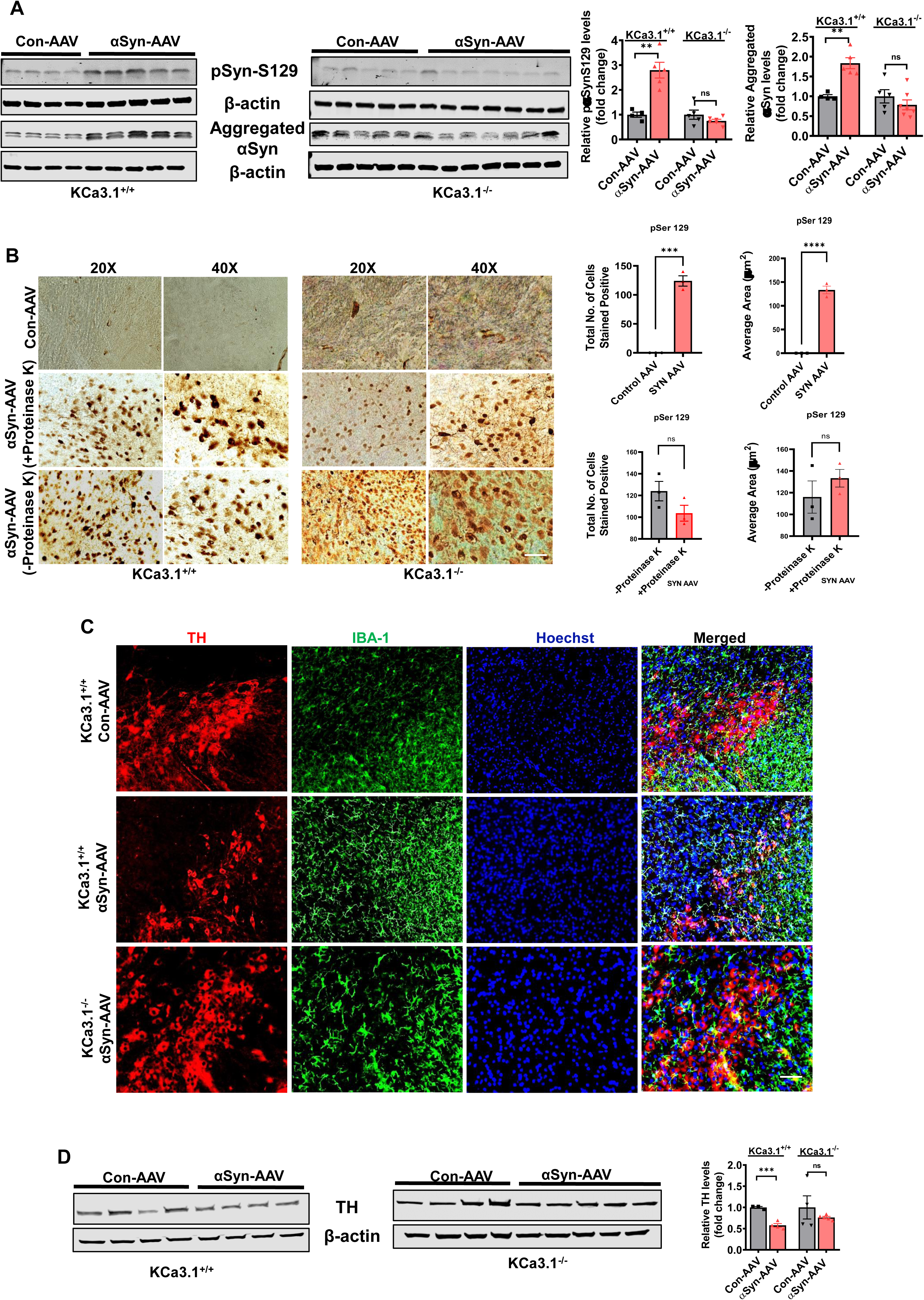
The effect of KCa3.1 deficiency on abnormal αSyn pathology in αSyn-AAV-inoculated mice. (A) Western blot analysis of striatal lysates at 6 wk post-αSyn-AAV inoculation normalized to β-actin showing KCa3.1 KO lowered aggregated and phosphorylated αSyn. (B-C) KCa3.1 gene deletion abolished αSyn-AAV-induced αSyn pathology at 6 wk post-viral injection. (B) Representative αSyn immunohistochemical images and quantification of pSer129-positive inclusions in the presence or absence of Proteinase K treatment showing KCa3.1 KO ameliorates αSyn-AAV-induced deposition of Proteinase K-resistant αSyn at 4 mo after inoculation. Scal bars: 50 µM. (C) Representative immunostaining for TH and IBA-1 of SNpc dopaminergic neurons. (D) Immunoblot blot analysis of TH using an antibody against TH. The relative band intensities were normalized to β-actin. Data represented as mean ± SEM from 4-6 per group. Significance was determined by one-way ANOVA followed by a Tukey’s post hoc test. *p≤0.05, **p<0.01, ***p<0.001.

Next, we sought to determine whether KCa3.1 KO influences αSyn-AAV-induced nigral DAergic neurodegeneration. Consistent with previous reports, at 16 wk post-αSyn-AAV inoculation, a marked decrease in the expression of TH was evidenced in the nigral brain region (Fig. 9C), as assessed via WB studies in line with previous studies [61, 64]. Likewise, our immunohistochemical studies revealed that the αSyn-AAV-induced reduction in TH+ DAergic neurons was also ameliorated in KCa3.1 KO mice, suggesting that KCa3.1 was sufficient to promote reactive microgliosis, αSyn pathology and nigral DAergic neurodegeneration (Fig. 9C-D). Herein we demonstrate that KCa3.1 in a central regulator of reactive microgliosis, a-syn pathology, and associated nigrostriatal DAergic neurodegeneration using an a-synculeinopathy mouse model of PD and that targeting KCa3.1 may represent a therapeutic strategy for PD and related a-synucleinopathies.

## DISCUSSION

KCa3.1 channels are highly expressed in activated microglia and are key mediators of both neurodegeneration and neuroinflammation, which is well supported by genetic and pharmacological studies [27, 30, 50]. In this study, using postmortem PD brain sections, primary microglial cultures, and multiple PD mouse models, we provide compelling evidence that KCa3.1 is upregulated within reactive microglia, and that its activation and the induction of the Fyn/STAT1 signaling axis play essential roles in driving the proinflammatory reactive phenotype of microglia in the context of α-synucleinopathy and in response to MPTP and LPS. Pharmacological or genetic inhibition of KCa3.1 reduced the αSyn_Agg_-induced proinflammatory reactive phenotype in microglial cells in culture and decreased neuroinflammation, PD-like neuropathology and associated neurobehavioral deficits in an MPTP and α-synucleinopathy mouse models of PD. KCa3.1 KO mice also ameliorated reactive microgliosis and brain cytokine levels following LPS challenge in vivo. Moreover, both transcriptomic and proteomic analyses further validated the proinflammatory profile of primary microglia stimulated with αSyn_Agg_, whereby ACOD1 upregulation was found to positively correlate with iNOS, IL-1β, and IL-6 generation. The great majority of inflammatory genes that were identified in the current study are consistent with previous studies demonstrating the upregulation of innate immune genes in α-synucleinopathy models [21, 37]. Collectively, our studies highlight the therapeutic utility of KCa3.1 inhibitors in reducing PD-pathogenesis and show that KCa3.1 is sufficient to promote neuroinflammation-associated nigral DAergic neurodegeneration presumably through a Fyn/STAT1 proinflammatory signaling mechanism in PD-vulnerable brain regions in experimental Parkinsonism as shown previously [35, 65].

Emerging evidence from our group and others has demonstrated the role for KCa3.1 in proinflammatory microglia polarization and neurodegeneration by triggering neuroinflammation in experimental models of neurodegeneration [25, 27, 30, 50, 66, 67]; however, the role of KCa3.1 in PD pathophysiology remains poorly understood. In the current study, we demonstrate that αSyn_Agg_ induced proinflammatory mediator generation and oxidative stress markers via elevated expression of KCa3.1 in both *in vitro* and *in vivo* models of PD. Lu et al. [67] found KCa3.1 pharmacological inhibition via senicapoc afforded marked DAergic neuroprotection via reduced generation of the proinflammatory cytokines TNF-α and IL-1β in MPTP-treated mice. Moreover, LC/MS analysis revealed sufficient target engagement in the cortical brain region of senicapoc-treated mitopark (MP) mice. Likewise, the anti-inflammatory effects of senicapoc were replicated, using another KCa3.1 inhibitor, TRAM-34, which diminished the generation of pro-inflammatory cytokines such as IL-1β and IL-6 in the sera of αSyn_PFF_ mice. Furthermore, TRAM-34 was found to abolish KCa3.1 channel activation in αSyn_Agg_-treated primary microglial cells. This is significant because this response positively correlated with reduced microgliosis, which is a significant driver of nigral DAergic neurodegeneration [68]. The TRAM-34 displayed a trend toward preservation of nigral DAergic neuronal integrity and marked protection against neurobehavioral deficits following α-synucleinopathy. TRAM-34 also has been shown to have similar effects in vivo where it was found to reduce brain infarction but also improve sensory motor deficits in mice subjected to MCAO [66]. Moreover, we previously demonstrated that KCa3.1 inhibition targets microglia, macrophages, and infiltrating T cells in an cerebral ischemia model [50]. Given that KCa3.1 is expressed in T cells and that KCa3.1 blockers can inhibit T cell function [30, 50], it is plausible that KCa3.1 inhibitors reduce T cell infiltration into the brains of α-synucleinopathy mice. Our studies highlight the therapeutic utility of KCa3.1 pharmacological blockers in not only ameliorating inflammation but also partially restoring the functionality of nigrostriatal DAergic neurons in diverse models of PD, and that it may be beneficial in the treatment of PD and related α-synucleinopathies. Thus, it remains plausible that a portion of the therapeutic effects of KCa3.1 inhibitor following systemic administration of the drug could be attributed to the inhibition of KCa3.1 in peripheral immune cells in the proposed mouse model of PD [69]. Thus, further studies are warranted to address the contribution of T cell modulation to the therapeutic effects of KCa3.1 inhibition in PD mouse models.

In the current study we performed a systematic investigation of proteomic changes associated with αSyn_Agg_-induced MMC microglial cell activation in the presence of TRAM-34. Our results demonstrate that out of the 24 mutually regulated proteins with STRING, three biological processes, including ncRNA processing, rRNA processing, and regulation of TLR4 pathway, were significantly altered (Fig. S1E-I). For example, ACOD1 and CD14 both are implicated in TLR4 signaling, with CD14 functioning as a positive regulator and ACOD1 as a negative regulator [40, 41]. Likewise, our RNA-seq analysis of microglia revealed that αSyn_Agg_ increased the expression of inflammatory immunomodulators such as ACOD1, which is linked to itaconate production, macrophage polarization, inflammasome activation, and oxidative stress [70], as well as iNOS, IL-6, IL-1β, CXCL-10 generation in PMG, while senicapoc pretreatment was found to downregulate the aforementioned inflammatory mediator generation, supporting the potential role of KCa3.1 in reactive microgliosis associated with α-synucleinopathy. Conversely, the expression of EGR3, IL-2R, and CD93 was upregulated in senicapoc/αSyn_Agg_-treated cells as compared to αSyn_Agg_-stimulated microglial cells, highlighting the involvement of genes corresponding to synaptic transmission, metabolism, survival, and immune response in PD-associated neuroinflammation. Together, our studies highlight the central role of KCa3.1-driven neuroinflammation in α-synucleinopathy and MPTP induced neurodegeneration further emphasizing the relevance of studying the αSyn_Agg_-induced microglial activation response to better understand PD pathology. However, a more comprehensive characterization of inflammatory mediators in a mouse model of synucleinopathy is needed to better understand PD pathogenesis.

Disease-associated microglia (DAM) has been shown to exhibit higher levels of STAT1, TLR2, and Fyn kinase and skews microglial polarization toward a proinflammatory phenotype and resultant nigral DAergic neurodegeneration in diverse inflammatory disease models [48, 71–73]. We observed increased Fyn expression and activation and its interaction with KCa3.1 as assessed via proximity labeling in αSyn_Agg_-stimulated microglial cells, indicating that Fyn-mediated KCa3.1 upregulation supports the proinflammatory response after α-synucleinopathy. In line with the detrimental role of Fyn signaling in neuroinflammation in the context of α-synucleinopathy, inhibition of Fyn via gene silencing or genetic knockout was associated with reduced proinflammatory mediator generation and downregulation of KCa3.1 expression. For example, our mechanistic studies revealed that Fyn is an upstream regulator of KCa3.1 upregulation in response to treatment with αSyn_Agg_, whereby Fyn genetic KO was found to reduce αSyn_Agg_-induced KCa3.1 expression in primary microglia, which is consistent with previous studies demonstrating the anti-inflammatory effects associated with Fyn inhibition in α-synucleinopathic mice [18, 21, 74–76]. Additionally, we also found that pharmacological inhibition and genetic depletion of Fyn reduced STAT1 activation in response to LPS or αSyn_Agg_, respectively. Our studies also reveal that Fyn is a critical regulator of STAT1 activation, which functions within macrophages and other innate immune cells to promote a proinflammatory phenotype in response to IFN-γ treatment [77–79]. The phosphorylation of STAT1 at tyrosine 701 (Y701) and serine 727 (S727) leads to the formation of STAT1 homodimers and their subsequent translocation to the nucleus in response to interferon signaling [80]. As evidenced in our study it is plausible that αSyn_PFF_-induced IFN gamma release may trigger STAT1 activation in a FYN dependent manner. Conversely, treatment of cells with fludarabine, a STAT1 inhibitor decreased STAT1 activation, with an accompanying reduction in kca3.1 expression and associated generation of proinflammatory cytokines in response to stimulation of microglial cells in culture with αSyn_PFF_. Our results support a novel signaling mechanism by which microglia fyn triggers stat1 activation which in turn binds to the promoter region of kca3.1 leading to channel upegulation and activation of neuroinflammation associated nigral daergic neurodegeneration via a feed back mechanism that sustains each other namely FYN/STAT1 signaling axis and kca3.1 eventually leading to an exaggerated neuroinflammation associated nigral daergic neurodegeneration-in response to alpha synucleinopathy.

It is noteworthy that microglia-specific Fyn deficiency ameliorates the microglial reactive phenotype and generation of proinflammatory mediators such as TNF-α and IL-1β in an LPS neuroinflammation mouse model of PD. Moreover, we previously demonstrated that Fyn KO mice afford DAergic neuroprotection by ameliorating neuroinflammation in an αSyn-AAV mouse model of PD [21]. Because Fyn activation is observed in multiple neurodegenerative disease models such as AD and PD [48, 81, 82], it is likely that Fyn inhibition may be associated with multifaceted therapeutic effects that are linked via the inhibition of multiple proinflammatory signaling cascades and subsequent preservation of nigral DAergic neuronal integrity in experimental Parkinsonism. We speculate that increased expression of KCa3.1 in an Fyn/STAT1-dependent manner results in an amplified proinflammatory microglial phenotype, thereby leading to the pronounced loss of nigral DAergic neurons in experimental Parkinsonism. Our results identify a novel signaling mechanism by which the Fyn/STAT1 signaling axis promotes KCa3.1 activation in microglia, thereby sustaining each other and eventually leading to pronounced neuroinflammation-associated nigral DAergic neurodegeneration in response to α-synucleinopathy.The effects of KCa3.1 deletion on neuroinflammation are reminiscent of Lu et al.’s [67] observations demonstrating diminished reactive microgliosis in MPTP-treated mice. In the current study, we employed the αSyn-AAV mouse model that has been routinely used to study the relationship between αSyn pathology and nigral DAergic neurodegeneration [83, 84]. Moreover, we validated our pharmacological and in vitro findings in an αSyn-AAV mouse model of PD using conventional KCa3.1 KO mice. First, a positive correlation between KCa3.1 gene ablation and reduced microglial reactive neurotoxic phenotype was evidenced in αSyn-AAV-inoculated mice, wherein morphometric characterization of microglia revealed that KCa3.1 KO resulted in a considerable reduction in several cellular parameters, such as cellular area, branch complexity, branch length, and number of branches. Moreover, KCa3.1 KO mice exhibited reduced generation of proinflammatory mediators and chemo-attractants, suggesting dysregulation of the innate and adaptive immune responses in the context of α-synucelinopathy. The fact that genetic deficiency of KCa3.1 is associated with reduced αSyn pathology and Fyn/STAT1 signaling events suggests that αSyn pathology may contribute to abnormal microglial function and nigral DAergic neuronal loss via the induction of the proinflammatory signaling cascade in proteinopathies such as PD. Collectively, these findings provide a framework for an improved understanding regarding the link between KCa3.1-mediated neuroinflammation and associated nigral DAergic neurodegeneration and Fyn/STAT1 signaling and reactive microgliosis as well as αSyn pathology in KCa3.1 KO mice following α-synucleinopathy. Additionally, studies using microglia-specific KCa3.1 KO mice may reveal the exact contribution of microglial KCa3.1 to α-synucleinopathy.

Although our studies provide novel insights into the mechanisms underlying KCa3.1-mediated neuroinflammation and nigral DAergic neurodegeneration and highlight the therapeutic potential of KCa3.1 inhibition in the treatment of PD, certain limitations need to be addressed in future studies. First, global KO of KCa3.1 may impact the signaling of other cell types besides microglia. Future studies aimed at utilizing microglia-specific KCa3.1 KO mice are necessary to determine whether microglial KCa3.1 is essential for the observed effect or whether it contributes to the activation of other immune cells such as astrocytes and T cells. Moreover, additional studies aimed at understanding the cross-talk between astrocytes and microglia in driving KCa3.1-mediated neuroinflammation-associated neurodegeneration are needed. Addressing these limitations will shed light on the critical role of KCa3.1-regulated mechanisms in neuroinflammation-associated nigral DAergic neurodegeneration.

In summary, our study describes a key regulatory role of KCa3.1 in driving pathological alterations in multiple PD models using diverse approaches. Our work demonstrates that neuroinflammation-associated nigral DAergic neurodegeneration is positively correlated with αSyn pathology in an α-synucleinopathy mouse model. We propose a model (Fig. 10) in which the pathological changes in microglia may occur via a Fyn driven STAT1 signaling mechanism in a KCa3.1-dependent manner, in response to αSyn_Agg_, which are reflected in an amoeboid reactive microglial phenotype and elevated neuroinflammation. Thus, our studies raise the possibility that KCa3.1 activation and the accompanying Fyn/STAT1 signaling axis may exacerbate neuroinflammation and subsequent PD-like neuropathology. Finally, our studies highlight the therapeutic benefits of KCa3.1-targeted pharmacological inhibitors in treating PD including α-syuncleinopathies.

**Fig 10.**
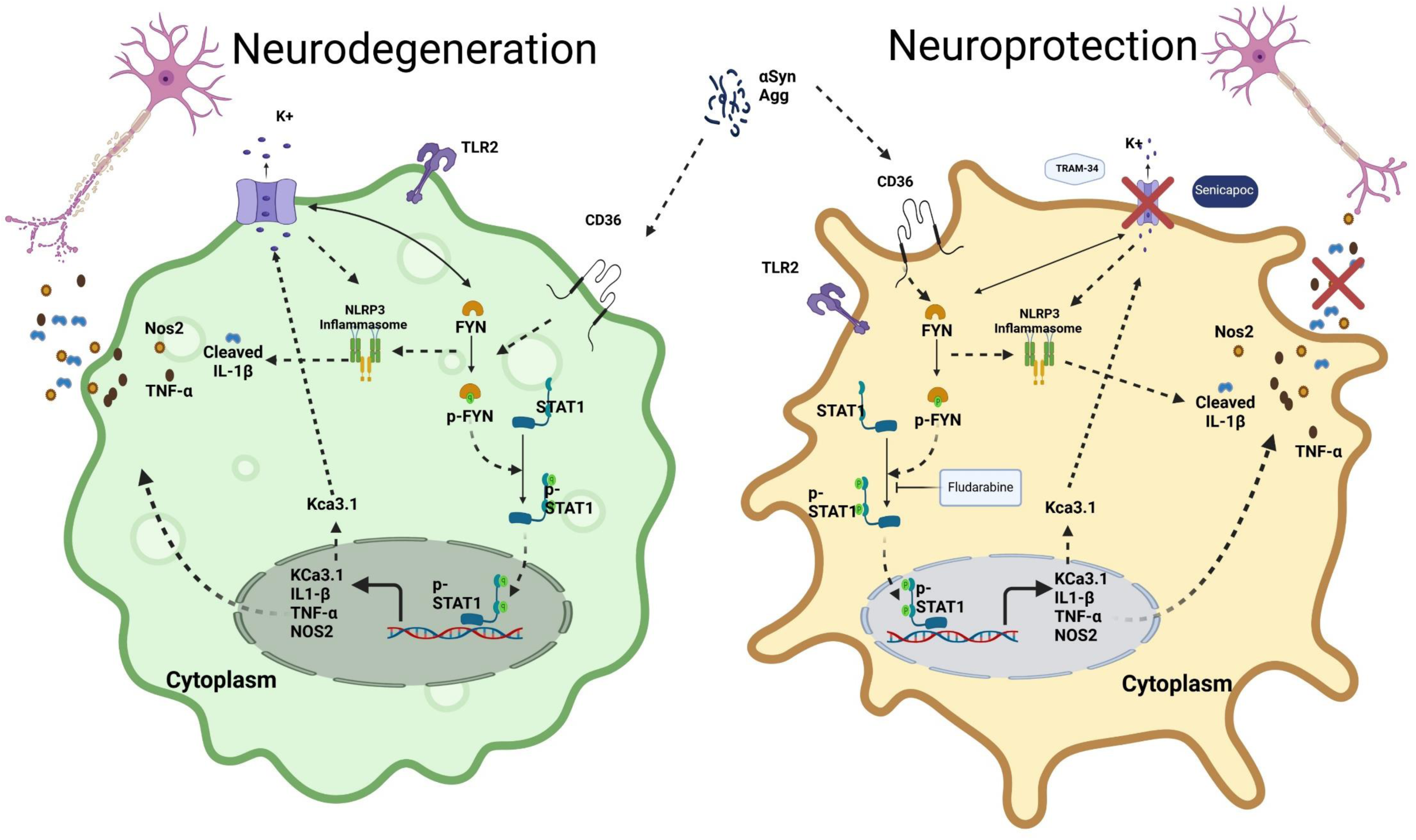
Aggregated αSyn-mediated NLRP3 inflammasome and pro-inflammatory cytokine activation pathway via a KCa3.1-dependent mechanism. Schematic representation of the involvement of KCa3.1 in microglia-driven neuroinflammation-associated nigral DAergic neurodegeneration in α-synucleinopathy. Diverse PD neurotoxins, including MPTP, LPS, and αSyn_Agg_, trigger a reactive microglial phenotype via a KCa3.1-dependent mechanism in an Fyn/STAT1 signaling cascade-dependent manner, thereby leading to the generation of proinflammatory cytokines, NLRP3 inflammasome activation markers, and oxidative stress mediators. A dysfunctional cross-talk between reactive microglia and dying DAergic neurons promotes α-synucleinopathy via a vicious self-propelling cycle that amplifies KCa3.1 activation in an Fyn/STAT1 signaling mechanism-dependent manner, eventually leading to nigral DAergic neurodegeneration. Both genetic depletion and pharmacological inhibition of KCa3.1 and downstream mediators were found to mitigate neuroinflammation and subsequent PD-like neuropathology. Increased expression of KCa3.1 in PD brains suggests that similar mechanisms operate in PD.

## Methods

### Human αSyn purification and aggregation

BL21 (DE3) cells transformed with a pT7–7 plasmid encoding WT human (h) αSyn were cultured from stocks in Luria broth medium containing 100 μg/mL kanamycin and incubated overnight at 37°C with shaking (pre-culture). The following day, this pre-culture was used to inoculate 1 L of Luria broth/kanamycin medium. Once the OD600 of the culture reached 0.6, 1 mM isopropyl β-D-1-thiogalactopyranoside (Invitrogen) was added to induce protein expression and incubated further at 37°C for 3 h before harvesting cells by centrifugation. Bacterial cell lysate was prepared by resuspending the cell pellet in approximately 30 mL of 500 mM NaCl, 50 mM Tris HCl at pH 7.6 using an Omni homogenizer. The lysate was then filtered using a 0.2-µm syringe filter and dialyzed overnight with 3,500-kDa dialysis tubing in 20 mM Tris HCl buffer at pH 8.0. The dialyzed lysate was collected and centrifuged at 15,000 × g for 10 min, and the lysate filtered using a 0.2-μm syringe filter and applied to a GE Sephacryl S-200-HP column using the AKTA Pure FPLC unit. The αSyn-containing fractions were identified by SDS-PAGE with Coomassie blue staining and pooled. The pooled fractions were dialyzed overnight against 20 mM Tris HCl at pH 8.0. Protein concentration was determined via a NanoDrop 2000 Spectrophotometer and aliquoted to 1 mg/vial and stored at −80°C for further applications.

### Transfections

The pre-designed STAT1 and scrambled siRNAs were purchased from Santa Cruz. A mouse microglia cell (MMC) line was used in the siRNA transfections with Lipofectamine 3000 reagent according to the manufacturer’s protocol. The cells were plated at 2 × 10^6^ cells/well in 6-well plates one day before transfection. For each well, 1 nM of STAT1 siRNA or an equal amount of scrambled siRNA was mixed with 5 µL of Lipofectamine 3000 and added to the cells. Then, 48 h after the initial transfection, the cells were treated with αSyn_Agg_ for 24 h. RT-qPCR was performed to verify the success of the transfections.

### Cell Culture

#### Mouse microglia cell (MMC) line

The MMC line (kind gift from Dr. D. Golenbock, University of Massachusetts) was derived by viral transduction of primary microglia. The maintenance media for the MMCs was DMEM-F12 containing 10% FBS, 1% penicillin, streptomycin, glutamine, and sodium pyruvate, while the experiments were conducted in the same media with reduced FBS (2%). Cell cultures were maintained in a humidified incubator at 37°C and 5% CO_2_.

#### Primary glial culture

Postnatal day 1 mouse pups were used to prepare primary microglial cultures based on a slightly modified technique described by Sarkar et al. [85]. The brains were harvested, and their meninges were removed and then placed in DMEM/F12 supplemented with 10% heat-inactivated FBS, 1% penicillin, 1% streptomycin, 1% L-glutamine, 1% nonessential amino acids, and 1% sodium pyruvate. The harvested brain tissues were immersed in 0.25% trypsin-EDTA and gently agitated for 15 min. The brain tissues were washed three times to stop trypsin activity and then triturated gently to prepare a single-cell suspension. The suspension passed through a 70-µm nylon mesh cell strainer to remove debris. The cell suspension was then made up in DMEM/F12 complete medium and seeded into T-75 flasks, which were incubated in a humidified 5% CO_2_ atmosphere at 37°C. We used a CD11b magnetic-bead separation kit to purify microglial cells from confluent mixed glial cultures and achieved 97% purity. Then the separated microglial cells were placed on the plate and allowed 48 h for recovery. The positive fraction containing microglia was plated on T-75 flasks containing the growth medium. Cells were allowed to grow overnight in growth medium in a 37°C incubator, and the next day the microglia were split and used for different experiments. Upon isolation of microglia using the magnetic bead technology, the negative fraction, predominantly consisting of astrocytes, was plated in T-75 flasks containing the growth medium.

### Human samples

SN tissues from patients with DLB and age-matched controls were obtained from the Miami Brain Endowment Bank at the University of Miami (Miami, Florida, USA). The average age of the patients with DLB was 80 ± 7 y, and the unaffected controls were aged 80 ± 6 y. The samples were processed and analyzed blindly.

### Animal studies, treatments and stereotaxic surgery

We purchased 8- to 12-wk-old male and female C57BL/6J mice from Charles River Labs (Wilmington, MA). KCa3.1 KO mice were generated and kindly provided by Dr. Heike Wulff in her laboratory at University of California (UC), Davis. MitoPark mice were generated by Dr. Nils-Goran Larsson and colleagues at the Max Planck Institute for Biology of Ageing by conditionally knocking out TFAM in cells expressing the dopamine transporter (DAT), as described in Ekstrand et al. [86]. The colony was kindly provided by their lab. The Fyn^−/−^ mouse colony was originally obtained from Dr. Dorit Ron’s laboratory at the UC, San Francisco. The Fyn^−/−^ used in these studies were bred at the Iowa State University (ISU) animal care facility and are also available from Jackson Laboratory (stock number 002271). Fyn cKO mice were generated at the ISU animal facility using cre/loxp technology. Briefly, tamoxifen conditional knockouts were generated by first crossing *Fyn*^(LacZ/0)^ with *Flp*^(Tg/0)^ mice to generate the floxed *Fyn* allele, denoted as *Fyn*(f). Next, these mice were bred with *Cx3cr1-GFP-Cre*^(Tg/Tg)^ mice to generate the heterozygotes *Fyn*^(fl/+)^:*Cx3cr1-GFP-Cre*^(Tg/0)^. Finally, the heterozygous offspring were bred together to acquire the Fyn cKO (*Fyn*^(fl/fl)^:*Cx3cr1-GFP-Cre*^(Tg/0)^) mice. All animals were allowed to acclimate to the new environment for 1 wk and were maintained in an environment with a temperature of 22 ± 2°C and 50 ± 10% humidity under a 12-h light-dark cycle with *ad libitum* access to food and water. All animal experiments were approved by the Institutional Animal Care and Use Committees of both ISU and the University of Georgia.

### Senicapoc studies in the MPTP mouse model

Given that men are at a higher risk of developing PD than women [87], we utilized male mice in these studies. Male C57BL/6 mice, aged 10–12 wk and weighing 24–28 g, were housed under standard conditions as described above. Mice were randomly assigned into four groups (n=8 per group) and subsequently allocated for either WB or immunofluorescence analyses: 1) vehicle control (Miglyol-812 (caprylic/capric triglyceride, Spectrum Chemicals), 2) MPTP (25 mg/kg, i.p. for 5 d), (3) senicapoc (50 mg/kg, i.p.), and (4) MPTP+senicapoc (50 mg, i.p). Senicapoc was administered twice daily for 11 d. One day before MPTP treatment, senicapoc was administered at 12-h intervals for a total of 2 doses. Thereafter, it was administered once before and once after MPTP treatment each day for a total of 5 d and continued for another 6 d until the day of sacrifice (11 d post-MPTP treatment). At the end of the treatment period, brains were harvested for immunohistochemistry or WB analyses.

### TRAM-34 studies in the αSyn_Agg_ mouse model

Male C57BL/6 mice (6- to 8-wk-old) were housed under standard conditions (22°C, 30% relative humidity, a 12-h light cycle, *ad libitum* food and water). Mice were randomly divided into two groups (control or αSyn_Agg_). An intrastriatal αSyn_Agg_-induced neuroinflammation model was used for this study as described by Panicker et al. [21]. The stereotaxic instrument was used with a 10-μL Hamilton syringe to inject 3 µL of αSyn_Agg_ directly into the striatum at the following stereotaxic coordinates in relation to bregma (mm): −2 ML, 0.5 AP, −4 DV. One subset of these animals received TRAM-34 (10 mg/kg, i.p.) daily starting at 4 mo and was sacrificed 1 mo later.

### The administration of h-αSyn-AAV into KCa3.1 WT (C57BL/6J) and KO mouse models

We used an adeno-associated virus vector, scAAV2/5-CBh-hu WT αSyn-WPRE3-enSV40pA, in which the expression of a wild-type (WT) h-αSyn gene was driven by a CBh promotor and enhanced by a woodchuck hepatitis virus post-transcriptional regulatory element (WPRE) and encoded with a polyadenylation sequence. The AAV vector was purchased from Charles River Labs (Cat. No: GD1006-RV), and the genome copy (GC) titer used in the injections was 3.34×10^12^ GC/mL. Mice were randomly divided into 4 groups (n = 4 – 6 per group); Group 1: KCa3.1 WT + Con-AAV (AAV 1/2 Null Empty, #GD1004-RV, Charles River Labs), Group 2: KCa3.1 WT + αSyn-AAV, Group 3: KCa3.1 KO + Con-AAV, Group 4: KCa3.1 KO + αSyn-AAV. For stereotaxic surgeries, mice were anesthetized using isoflurane. A digital stereotaxic instrument was used with a 10-µL Hamilton syringe to deliver either Con-AAV or αSyn-AAV (2 µL) into the right dorsal SN of isoflurane-anesthetized KCa3.1 WT and KO mice at the following stereotaxic coordinates relative to bregma (mm): −3.1 AP; +1.2 ML and −4.6 DV. After drilling a 1-mm burr hole in the skull, a 2-μl volume of solution was infused at the target site at the rate of 0.2 µL per minute. The needle was held in place for at least 5 min after injection to minimize retrograde flow along the needle tract. Mice were given a subcutaneous injection of sterile lactated Ringer’s solution to facilitate recovery and were placed on a heat-pad until complete recovery from anesthesia.

### LPS injections in the KCa3.1 WT and KO mouse models

The nigral neuroinflammatory response was also studied using the systemic LPS injection model [48], which induces chronic neuroinflammation and progressive DAergic degeneration in mice. KCa3.1 WT and KO mice received a single injection of LPS (5 mg/kg, i.p.) used to induce neuroinflammation in the brain and were sacrificed 20 h later.

### Isolation of adult microglia from the mouse brain

Adult microglial cells were isolated from LPS-injected brains using a modified protocol of Bordt *et al*. (2020). Briefly, after perfusion with ice-cold PBS, brains were dissected, and enzymatically digested using the Adult Brain Dissociation Kit (Miltenyi Biotec, Germany) for 30 min at 37°C in a heated gentleMACS Octo Dissociator. Further processing was performed at 4°C. Tissue debris was removed by passing the cell suspension through a 70-μm cell strainer. Myelin was removed using a Percoll gradient centrifugation method. After myelin removal, cells were incubated with CD11b (Microglia) MicroBeads (Miltenyi Biotec, #130-093-634) in MACS buffer (PBS supplemented with 0.5% BSA and 2 mM EDTA) for 15 min in a refrigerator (2-8°C) and cells were washed by adding 1−2 mL of buffer and centrifuged at 300 × g for 10 min. Next, we aspirated the supernatant completely and resuspended the cells in 500 µL of MACS buffer. CD11b+ cells were separated in a magnetic field using LS columns (Miltenyi Biotec, # 130-042-401). The amounts of antibodies and magnetic beads were calculated based on the number of cells obtained after myelin removal, using the manufacturer’s guidelines. Both the CD11b+ and CD11b-(effluent) fractions were collected and used for further analyses. The isolated cells were resuspended in Trizol reagent for RNA isolation and applications, or in RIPA extraction for WB analysis.

### Motor function test

For open-field test, a VersaMax system (VersaMax monitor, model RXYZCM-16, and analyzer, model VMAUSB, AccuScan, Columbus, OH) was used for monitoring locomotor activity [88, 89]. The clear Plexiglas chamber has dimensions of 40 × 40 × 30.5 cm, and is covered with a ventilated Plexiglas lid. Infrared (IR) monitoring sensors are located every 2.54 cm along the full perimeter of the square chamber (16 IR beams along each side). For horizontal and vertical activity and corresponding plots, mice were acclimated for 3 d and trials were started at a later time point prior to recording for 10 min using the VersaMax system.

The RotaRod equipment (AccuScan) was used to test coordination of movement as described by Ghosh et al. [90]. Briefly, we used the constant speed profile to measure the motor activity in the animals. The time spent on rod rotating at 20 rpm was measured for a maximum of 20 min with three trials, each of which ended with a mouse falling from the rod. The mean time spent or latency to fall by the mice on the rod was measured and compared between the groups.

### Mass spectrometry proteomic analysis

MMCs were grown to 75% confluence and then exposed to αSyn_Agg_ (1 μM) or αSyn_Agg_ plus TRAM34 (1 µM) for 24 h. Cells were harvested and centrifuged into pellets. Next, the cell pellets were incubated in 300 μL of urea lysis buffer (8 M urea, 100 mM NaHPO4, pH 8.5), along with HALT protease and phosphatase inhibitor cocktail (1:100) (Pierce) for 30 min at 4°C. Then, the lysates were sonicated (Sonic Dismembrator, Fisher Scientific) three times for 5 s with 15-s intervals of rest at 30% amplitude to disrupt nucleic acids and vortex-mixed. Bicinchoninic acid (BCA) was used to quantify protein concentration [37]. Protein lysates were reduced with 1 mM dithiothreitol (DTT), followed by 5 mM iodoacetamide and lastly overnight digestion using trypsin/Lys-C. Formic acid was used to stop the digestion and then the samples were dried out using a SpeedVac. The samples were desalted using C18 columns (Nest Group BioPureSPN Mini, HUM S18V) before drying again in a SpeedVac. PRTC standard (Pierce part #88320) was spiked into the sample to serve as an internal control. The peptides were then separated by liquid chromatography and analyzed by MS/MS by fragmenting each peptide. The resulting intact and fragmentation pattern was compared to a theoretical fragmentation pattern from either MASCOT or Sequest HT for peptide and protein identification. The PRTC areas were used to normalize the data between samples. For liquid chromatography, we used the Thermo Scientific EASY nLC-1200 coupled to a Thermo Scientific Nanospray FlexIon source. For mass spectrometry, we used the Thermo Scientific Q Exactive Hybrid Quadrupole-Orbitrap Mass Spectrometer with an HCD fragmentation cell in conjunction with Proteome Discover 2.0. Volcano plots and heatmaps were generated using Graph Pad 8.0, Venn diagrams using Microsoft PowerPoint, and Cytoscape for Gene Ontology analysis and graphing of STRING-generated data.

### RNA sequencing and analysis

The primary microglial cells were cultured and isolated as described in previous publications [18, 85]. At the end of the treatment period, cells were pelleted and subsequently flash frozen and stored at −80°C until the day of the analyses. The RNA was isolated with Qiagen RNeasy® Micro Kit (Qiagen, Germany) according to the manufacturer’s protocol. Isolated RNA sample quality was measured by High Sensitivity RNA Tapestation (Agilent Technologies Inc., California, USA) and quantified by AccuBlue® Broad Range RNA Quantitation assay (Biotium, California, USA). Paramagnetic beads coupled with oligo d(T)25 are combined with total RNA to isolate poly(A)+ transcripts based on NEBNext® Poly(A) mRNA Magnetic Isolation Module manual (New England BioLabs Inc., Massachusetts, USA). Prior to first-strand synthesis, samples were randomly primed (5′ d(N6) 3′ [N=A, C, G,T]) and fragmented based on manufacturer’s recommendations. The first strand is synthesized with the Protoscript II Reverse Transcriptase with a longer extension period, approximately 40 min at 42⁰C. All remaining steps for library construction were used according to the NEBNext® Ultra™ II Directional RNA Library Prep Kit for Illumina® (New England BioLabs Inc., Massachusetts, USA). Final library quantity was assessed by Qubit 2.0 (ThermoFisher, Massachusetts, USA) and quality was assessed by TapeStation D1000 ScreenTape (Agilent Technologies Inc., California, USA). The final library size was about 430 bp with an insert size of about 300 bp. Illumina® 8-nt dual-indices were used. Equimolar pooling of libraries was performed based on QC values and sequenced on an Illumina® Novaseq platform (Illumina, California, USA) with a read-length configuration of 150 PE for 40M PE reads per sample (20 M in each direction) [91]. FastQC (version v0.12.1) was applied to check the quality of raw reads. Trimmomatic (version v0.39) was applied to cut adaptors and trim low-quality bases with the default setting. STAR Aligner (version 2.7.10b) was used to align the reads. The package of Picard tools (version 3.0.0) was applied to check the quality of mapping. StringTie (version 2.2.1) was used to assemble the RNA-seq alignments into potential transcripts. FeatureCounts (version 2.0.6) or HTSeq (version 2.0.3) was used to count mapped reads for genomic features such as genes, exons, promoters, gene bodies, genomic bins, and chromosomal locations [92]. DESeq2 (version 1.18.1) was used to process the differential analysis. Gene Ontology Analysis was done using the ClusterProfiler package in R [93].

### Immunoblotting

Protein lysates from cells and brain tissues were prepared in RIPA buffer (20 mM Tris-HCl, pH 7.4, 2.5 mM EDTA, 1% Triton X-100, 1% sodium deoxycholate, 1% SDS, 100 mM NaCl, 100 mM sodium fluoride) with protease and phosphatase inhibitors. For WB, equal amounts of protein lysates (30 μg) were run on a 10-12% SDS-PAGE gel and transferred to a nitrocellulose membrane. After blocking for 1 h, primary antibodies against IBA-1 (Abcam # ab178846), iNOS (Abcam # ab202417), Fyn (Invitrogen # MA1-19331), pFynY416 (Cell Signaling # 6943), TH (Sigma Millipore # AB1542), NLRP3 (Cell Signaling # 15101), αSyn aggregates (MJFR, Abcam # ab209538), pαSynS129 (Abcam # ab209422), KCa3.1 (Inivtrogen # PA5-106671), TGF-β (3711S), IL-10 (mAb #12163), CD68 (ab201340), Arg-1 (mAb #93668), STAT1 (#9172), pSTAT1 (#9177) and β-actin (Millipore, Billerica, MA, USA) were incubated at 4°C overnight. The membranes were then incubated with secondary antibodies (IR conjugated) at RT for one hour and images were captured via a LI-COR Odyssey imager. Densitometric analysis was done using ImageJ software.

### Immunofluorescence staining

Immunohistochemistry was performed in mouse as well as human nigral sections as described in our previous publications with slight modifications [90, 94]. Mice were transcardially perfused with 4% paraformaldehyde (PFA) and were cryoprotected with 30% sucrose the following day. Brains embedded in OCT at −80°C were cryosectioned to 30-µm sections, which were then stored in cryosolution (ethylene glycol and sucrose) until use. Citraconic anhydride was used for human samples and 10 mM citrate buffer at pH 7.4 was used for animal samples for antigen retrieval. The sections were washed with PBS following antigen retrieval, and blocked with a blocking buffer (2% BSA, 0.5% Triton X-100, and 0.05% Tween-20). After blocking, the sections were incubated in primary antibodies overnight at 4°C. Next day, sections were washed with PBS, and incubated in secondary antibodies for 1 h, and stained with Hoechst for nuclear visualization. In the end, sections were mounted on precoated slides and allowed to dry overnight before visualization under a fluorescence microscope.

For immunoperoxidase staining [75], mice were deeply anesthetized and perfused with saline (0.9%) and 4% ice-cold PFA. The brain samples were subsequently removed and stored in PFA solution for 72 h at 4°C, followed by immersion in 30% sucrose at 4°C for the next 72 h. The brains were fixed in optimal cutting temperature (OCT) compound (company indicates), and the midbrain region encompassing striatum and substantia nigra were serially cut into 30-µm sections using the cryo microtome (Leica cryostat CM3050, Heidelberg, Germany). The slide containing the sections was washed with PBS, incubated with methanol containing 3% H_2_O_2_ for 30 min, washed with 6x PBS for 5 min each, and blocked with 5% normal donkey serum, 0.5% Triton X-100 in PBS for 1 h at room temperature (RT). Then, the sections were incubated with an anti-TH antibody (1:1,500, mouse monoclonal) overnight at 4°C and washed in 3x PBS for 10 min each. Biotinylated secondary antibody was used for 1 h at RT, followed by incubation with an avidin peroxidase solution (ABC Vectastain Kit, Vector laboratories, Burlingame, CA) for 30 min at RT. Immunolabelling was observed using DAB (Diaminobenzidine) solution, which yields a brown stain, and images were captured using a Keyence BZ-0023 Microscope (Keyence, Osaka, Japan). The total number of TH^+^ neurons was counted using the Stereo Investigator software (MBF Bioscience, Williston, VT, USA). We used every sixth section of the substantia nigra [95], and they were visualized using an Eclipse TE2000-U (Nikon, Tokyo, Japan) inverted fluorescence microscope for stereological counting.

Immunofluorescence studies with primary mouse microglial cells were performed based on Ghosh et al. [95] with few modifications. The coverslips were coated with PDL and plated with approximately 20,000 cells. After the respective treatments, the cells were fixed with 4% PFA, washed in PBS, and blocked with buffer (PBS containing 2% BSA, 0.5% Triton X-100, and 0.05% Tween-20) for 1 h at RT. Cells on the coverslips were probed with the respective primary antibodies, specifically IBA-1 (1:1,000, goat polyclonal) and KCa3.1(1:500, rabbit polyclonal) diluted in PBS containing 1% BSA and incubated overnight at 4°C. Next day, cells on the coverslips were washed several times in PBS and incubated with Alexa Fluor 488 and Alexa Fluor 555 dye-conjugated secondary antibodies. Hoechst counterstain was used to stain nuclei, and the stained cells on the coverslips were mounted using Fluoromount medium on glass slides for visualization. Samples were visualized using an inverted fluorescence microscope and images were captured using Keyence BZ-X810.

### Multiplex cytokine Luminex immunoassays

We used the Bio-Rad Multiplex Cytokine Luminex kit for key proinflammatory cytokine assays. The assay was conducted according to the manufacturer’s protocol along with positive and negative controls with some modifications. After treatment, mice were sacrificed, and the desired brain regions were harvested and stored at −80°C. The tissue lysates were subsequently homogenized. For cell studies, whole-cell lysates were prepared. Briefly, 50 µL of cell lysate or tissue samples from each experimental group was incubated with multiplex beads conjugated with primary antibodies for 1 h at RT. After the incubation, the samples were washed and incubated with the detection antibody and biotin/streptavidin. Finally, the data were acquired using the Bio-Plex plate reader.

### Duo-link Proximal Ligation assay (PLA)

PLA was carried out according to our protocol described in Samidurai et al. [18]. Briefly, 10,000 primary microglial cells were seeded on PDL-coated coverslips in 96-well culture plates. After the treatments, the cells were washed with PBS and fixed using 4% PFA, blocked with blocking buffer (PBS containing 1.5% BSA, 0.5% Triton X-100, and 0.05% Tween-20), and probed with antibodies against Fyn and KCa3.1 and allowed to incubate at 4°C overnight. Next day, the positive and negative oligonucleotide (PLA probes) conjugated secondary antibodies were incubated with the cells. Following the secondary antibodies, the ligase was added to allow the oligonucleotides to hybridize to the PLA probes, and when close enough, these oligonucleotides form a closed circle. In the DNA circle, the antibody-conjugated DNA probes serve as a primer for amplification, and a DNA polymerase and nucleotides were added to generate repeated sequence product. Fluorescently labeled oligonucleotides allow visualization of the protein-protein interaction as single dots via fluorescence microscopy [96]. The protein-protein interactions appeared as green puncta, which were visualized and analyzed by a Keyence microscope (with 60X objective). We used ImageJ software to further quantify the PLA signals according to Gomes et al. [97].

### Whole-Cell Patch-Clamp

Whole-cell patch-clamp studies were carried out as described in Jin et al. [30]. Briefly, the conductance of a KCa3.1 channel was calculated based upon the slope of TRAM-34-sensitive KCa current: between −80 and −75 mV, whereby under the proposed experimental conditions involving an aspartate-based internal solution, the currents were primarily carried by the KCa3.1 channel uncontaminated by Kv1.3 (shows activation at voltages that are more positive than −40 mV), inward rectified K+ currents (which are evidenced at negative voltages > −80mV), or chloride solution.

### Pharmacokinetic studies for senicapoc

The basic pharmacokinetic properties (brain/plasma ratios) and oral bioavailability were assessed as described in Jin et al. [30]. Breifly, the brain concentrations of senicapoc were determined by LC-MS analysis using a Waters Acquity UPLC (Waters) interfaced to a TSQ quantum Access Max mass spectrometer (Thermo Fisher Scientific).

### ChIP study

Cell lysates were collected after the respective treatments. Working lysis buffer (1X) was added to the cell pellet, which was then resuspended by carefully pipetting up and down. The cell suspension was transferred to a fresh vial and incubated on ice for 10 min. The supernatant was carefully removed after the centrifugation. Working extraction buffer was added to the chromatin pellet and resuspended by carefully pipetting up and down, followed by sonication. After sonication, chromatin buffer was added to the samples at a 1:1 ratio and stored at −80°C. ChIP reactions were prepared by adding the STAT1 antibody (ab239360), chromatin, and other reagents to each well according to the manufacture’s instructions (ab156907). The reaction tubes were placed on a magnetic stand for 12 min to pelletize the beads to the sides of the wells. The pellets were washed several times, and the DNA was eluted by incubating with DNA-releasing buffer at 65°C for 15 min and then 95°C for 5 min in a thermal cycle. Purified DNA was used to analyze the KCa3.1 expression using qPCR. The qPCR reactions were conducted in an Applied Biosystems Quantstudio 5 Real-Time PCR System using SYBR Green/ROX qPCR Master Mix as described by Lee et al. [50].

### HPLC

The striatal tissue samples were prepared and quantified, as described previously [15, 16]. Prior to the analysis, the neurotransmitters from the tissues were extracted using an antioxidant extraction solution (0.1 M perchloric acid, containing 0.05% Na_2_EDTA and 0.1% Na_2_S_2_O_5_) and isoproterenol (as internal standard). Using a reversed-phase column with a flow rate of 0.6 mL/min, the DA and 3,4-dihydroxyphenyl-acetic acid (DOPAC) were separated isocratically. An HPLC system (ESA, Inc, Bedford, MA, USA) with an automatic sampler equipped with refrigerated temperature control (Model 542; ESA, Inc) was used for these experiments.

### RNA isolation and real-time RT-PCR

Total RNA was isolated from cells or substantia nigra with TRIzol (Thermo Fisher, Catalog # 15596026) according to the manufacturer’s protocol. RNA purity and concentration were assessed using a NanoDrop Spectrophotometer. RNA was reversely transcribed with the High-Capacity cDNA Reverse Transcription Kit (Thermo Fisher Scientific, Catalog# 4374967) with 2 µg of total RNA per 20-µL reaction. Reactions were carried out in an Applied Biosystems PCR machine; the conditions were as follows: 25°C for 10 min, 37°C for 120 min, 85°C for 5 min, with final storage at 4°C. RT-qPCR reactions were conducted in an Applied Biosystems Quantstudio 5 Real-Time PCR System using SYBR Green/ROX qPCR Master Mix as described by Lee *et al.* [50]. Data are presented as a ratio of mean threshold (Ct) target gene expression and the housekeeping gene β-actin. Fold changes were calculated by the ΔΔCt method. Primer sequences are as follows

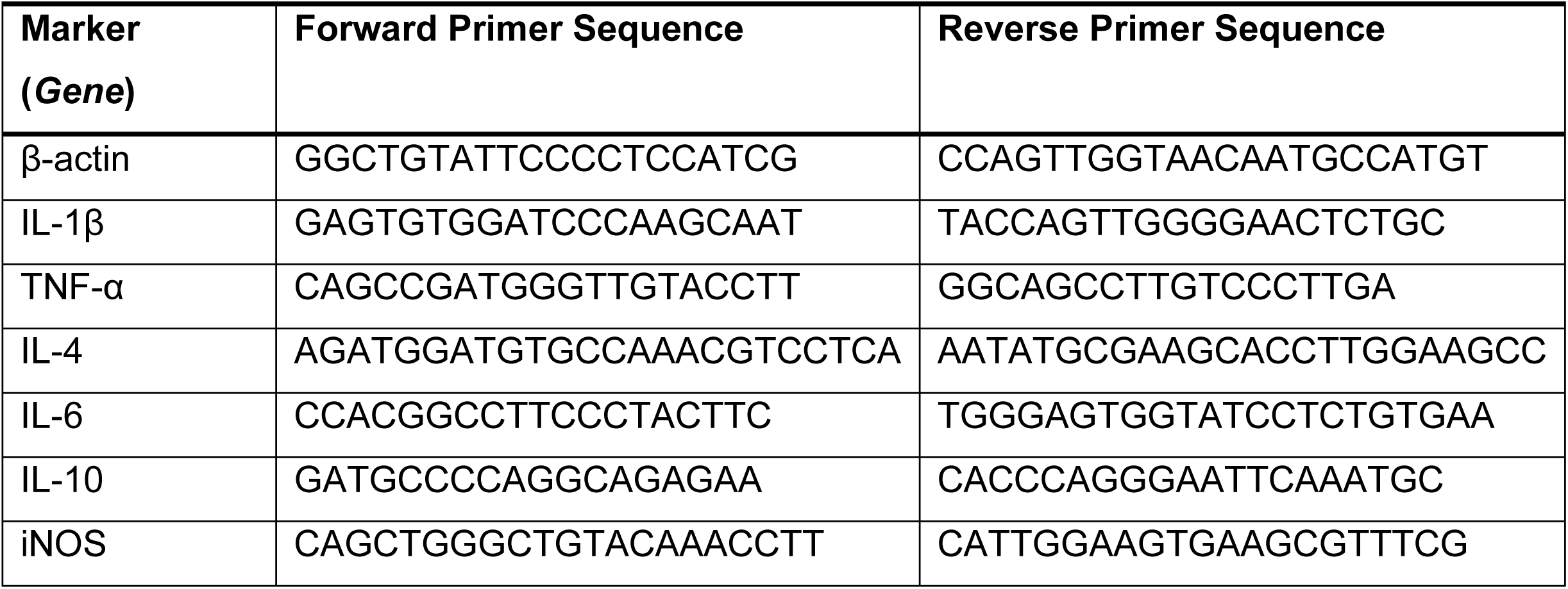

The CCL2, CCR2, KCa3.1 and Arg-1 primers were purchased from Qiagen.

### Morphological assessment of microglia cell counts

All analyses were performed using Image J software. Microglial morphometric analysis was carried out using skeletonized images to quantify microglia morphology in IHC images of the striatum and SNpc, following a modified method described in Putra et al. [98]. IBA-1-positive cells in the striatum and SNpc regions of brain tissue sections were imaged using a Keyence microscope that has a z-stack acquisition capability using a 20X objective or greater. Imaging parameters and software set-up were constant for all photomicrograph acquisitions in our experiments. To measure the intensity of the background staining, at least four images across each region of interest (ROI) were analyzed, with four non-stained regions identified based on knowledge of the antigen target. These regions were measured for pixel intensity, and the four values were averaged. Images were converted to 8-bit black and white, adjusted for contrast, and made binary with the ‘threshold’ plugin to remove background. The ‘despeckle’ plugin was applied to remove single pixels. Pixels were then enlarged by the ‘dilate’ plugin, making multiple single pixels become a cluster. The ‘outline’ and ‘fill holes’ plugins were then used to trace and fill in the pixelated structures, which were then processed by the ‘skeletonize’ plugin. Cell branches were then tagged with skeletal features, which included slab voxels representing process length, and endpoints. The cell body area of each microglia in each image was determined after converting the pixel into micrometer (μm) scale. The image was then binarized and skeletonized using ImageJ software. Data from each image (endpoints summed and branch lengths summed) were divided by the number of microglia somas in the corresponding image. The final data (endpoints/cell & branch length/cell) were entered into statistical software. Approximately 50–70 randomly selected microglia cells in the striatum and SNpc were analyzed in each group.

### Statistical analysis

Data were analyzed using a two-tailed t-test (two groups) or one-way ANOVA followed by Bonferroni’s post hoc analysis (PRISM 9.0 software, GraphPad, La Jolla, CA, USA). Data were represented as mean ± SEM. With Type I error set to α=0.05, we used p≤0.05 to determine significance for all statistical tests.

## Author Contributions

Conceptualization: M.S., K.C., H.W. and A.K. Investigation, Methodology, Data curation, Validation, Visualization, Formal analysis: M.S., K.C., S.S., E.M, H.N, L.S, A.K, A.E, C.J, B.N.P, and N.K. Project administration, Resources, Supervision, Validation, Funding acquisition: A.K., H.W, A.G.K., H.J. and V.A. Visualization, Writing - original draft, Writing - review & editing: M.S., K.C, H.J., G.Z., H.W, A.G.K and A.K.

## Data Availability Statement

All data needed to evaluate the conclusions in the paper are presented in the manuscript and Supplementary Materials. Any materials described in this manuscript may be obtained through a material transfer agreement.

## Acknowledgments

The authors would also like to thank Prashant Tarale for assisting in one of the qPCR experiments. We also acknowledge Admera Health for their contributions to the RNA sequencing and analysis. This study was supported by the National Institute of Neurological Disease and Stroke (grant number: R01 NS124226) to A.K. The CVM and UGA Foundation start grant and partial support from R01 NS121692 grant to A.G.K. are also acknowledged.

## Conflicts of Interest

A.G.K. has an equity interest in PK Biosciences Corporation and Probiome Therapeutics located in Athens, GA. The terms of this arrangement have been reviewed and approved by Iowa State University and the University of Georgia in accordance with their conflict-of-interest policies. Other authors declare no actual or potential competing financial interests.

## Notes

### Competing Interest Statement

The authors have declared no competing interest.

