## Supplementary figures 1-6 for "KCa3.1 Contributes to Neuroinflammation and Nigral Dopaminergic Neurodegeneration in Experimental models of Parkinson’s Disease"

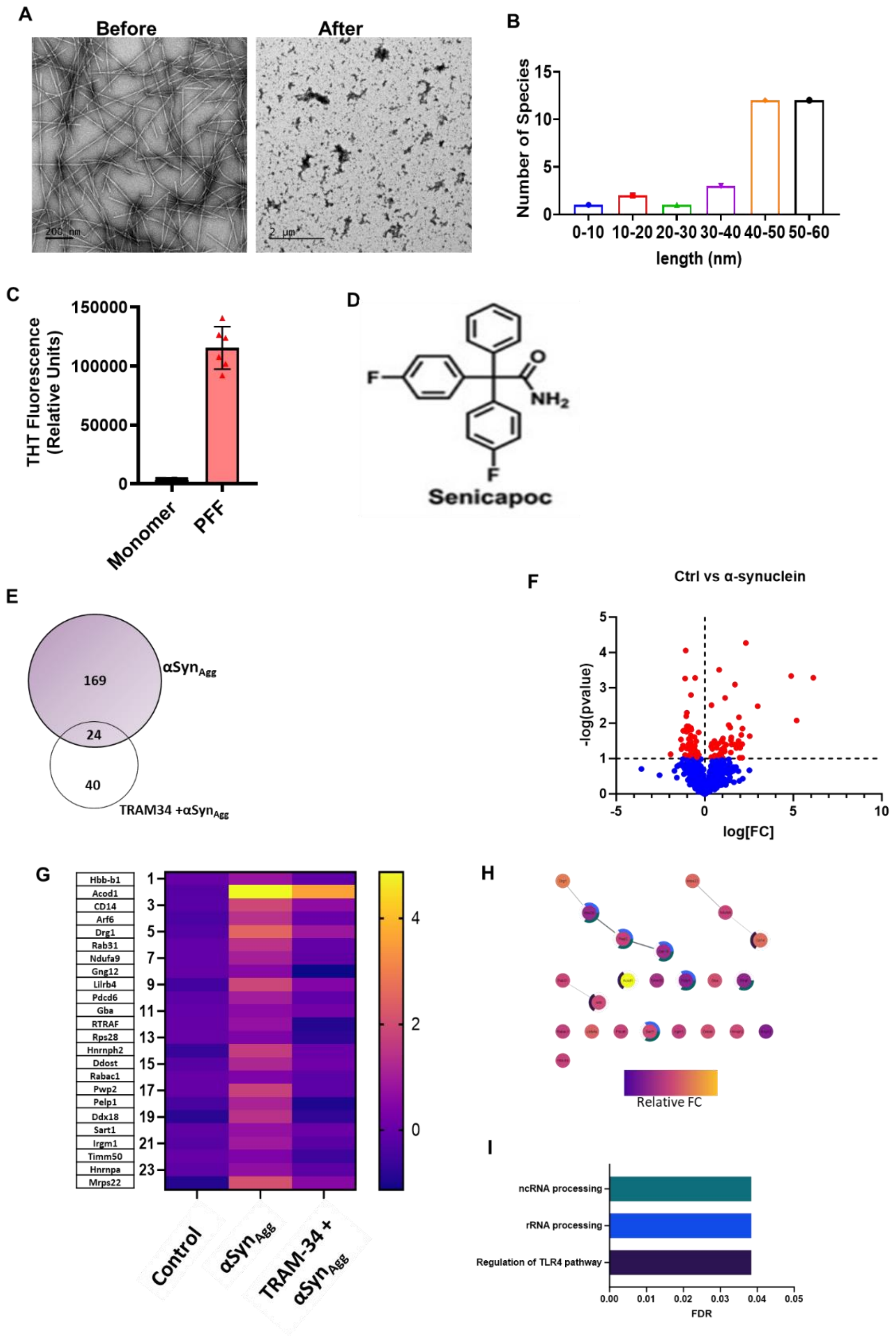


**Fig. S1.** (A) Representative transmission electron microscopic (TEM) images of $\alpha Syn_{Agg}$ before and after sonication. (B) Quantification of the size distribution of the $\alpha Syn_{Agg}$ after sonication acquired from the analysis of aggregates from the TEM images. (C) Results of Thioflavin T aggregation assay shown as ΔRFU value. (D) Chemical structure of senicapoc. (E) Volcano plot demonstrating differential protein levels of microglia treated with $\alpha Syn_{Agg}$ (1 µg/mL) when compared to control microglia. The cutoff was $p<0.1$, where red dots indicate significant changes, while blue indicates insignificant changes. (F) Venn diagram showing the total number of differentially changed proteins between the denoted groups: 169 for controls vs αSyn$_{Agg}$, 40 for αSyn$_{Agg}$ vs TRAM-34 + αSyn$_{Agg}$, and 24 shared proteins between the two comparisons. The cutoff was $p<0.1$. (G) Heatmap showing fold change values for the 24 shared proteins, demonstrating a general increase of these proteins by αSyn$_{Agg}$ and an attenuation by TRAM-34 inhibition. (H) STRING analysis of the 24 shared proteins revealing regulation of TLR4 pathway, ncRNA processing, and rRNA processing, as the top biological processes, with FDRs generated by STRING based on the number of identified proteins compared to the background total of proteins within those biological processes. Colored arches denote the protein's involvement in those biological processes: teal for ncRNA processing, blue for rRNA processing, and purple for regulation of TLR4 pathway. Data are mean ± SEM; n=3-6. *$p\leq0.05$, determined by one-way ANOVA followed by Dunnett's multiple comparison test.

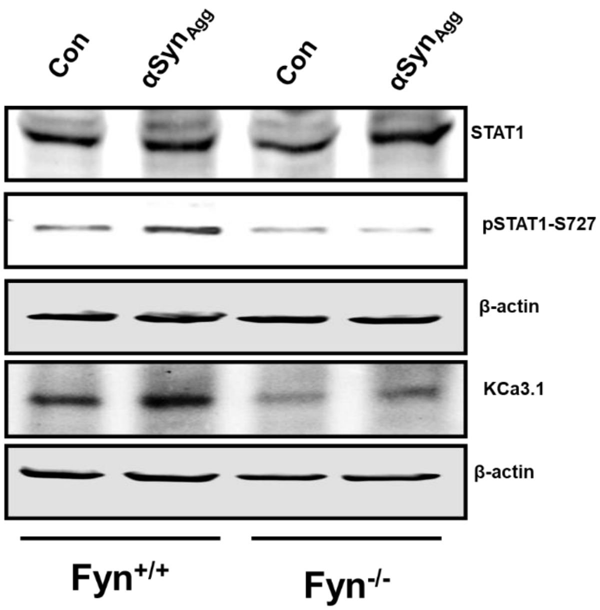


**Fig S2.** Fyn deficiency attenuates $\alpha Syn_{Agg}$-induced KCa3.1 expression and STAT1 phosphorylation in primary microglial cultures. Representative Western blot results showing reduced KCa3.1 and STAT1 phosphorylation in Fyn KO primary microglial cultures.

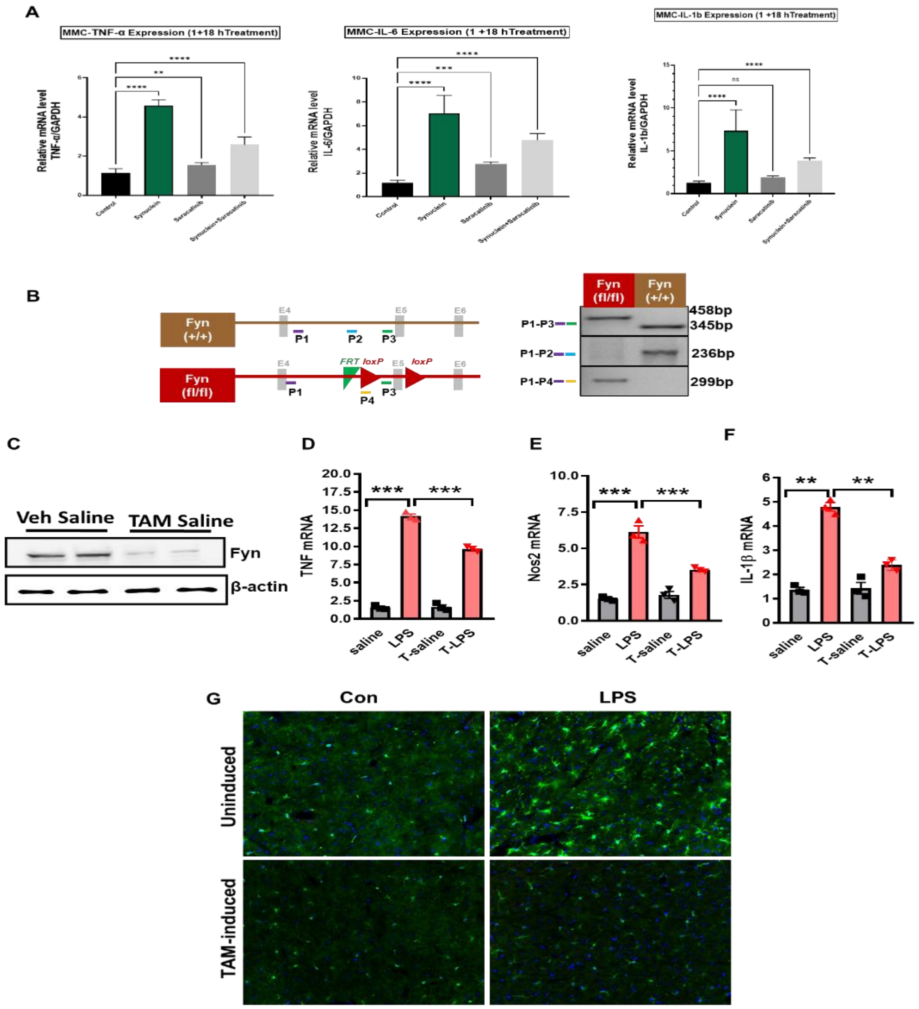

**Fig S3.** (A) RT-qPCR analysis of microglia treated with 1 µM αSyn$_{Agg}$ in the presence or absence of the Fyn kinase inhibitor saracatinib for 24 h revealing that saracatinib co-treatment inhibited αSyn$_{Agg}$-stimulated mRNA expression of TNF-α, IL-1β and IL-6. (B) Genotyping of Fyn$^{(fl/fl)}$/Cx3cr1-GFP-Cre $^{(Tg/0)}$ mice: Genomic DNA was isolated from the Fyn$^{(fl/fl)}$/Cx3cr1-GFP-Cre $^{(Tg/0)}$ mice generated in the breeding colony at ISU animal facility using the Qiagen Genomic DNA Isolation kit. Genomic DNAs underwent three independent PCR reactions using three primer sets, P1 & P2, P1 & P3, and P1 & P4. The presence of 458 bp (P1-P3) and 299 bp (P1-P4) confirms the presence of transgenic Fyn (fl/fl) floxed mice, while 345 bp (P1-P3) and 236 bp (P1-P2) confirm the presence of Fyn$^{+/+}$ mice. (C) Representative immunoblots showing scant expression of Fyn in isolated adult microglia from Fyn cKO mice as compared to Fyn$^{fl/fl}$ mice (n=4 per group). (D-F) Reduced mRNA expression of the proinflammatory markers TNF-α, iNOS, and IL-β in response to LPS treatment of microglia-specific Fyn knockout mice. (G) Diminished IBA-1 immunofluorescence staining in Fyn cKO microglia in the nigra of LPS-treated mice.

A

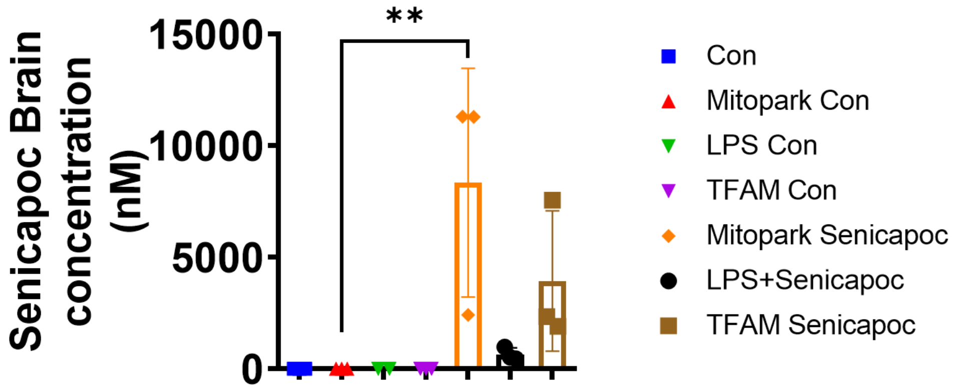


B

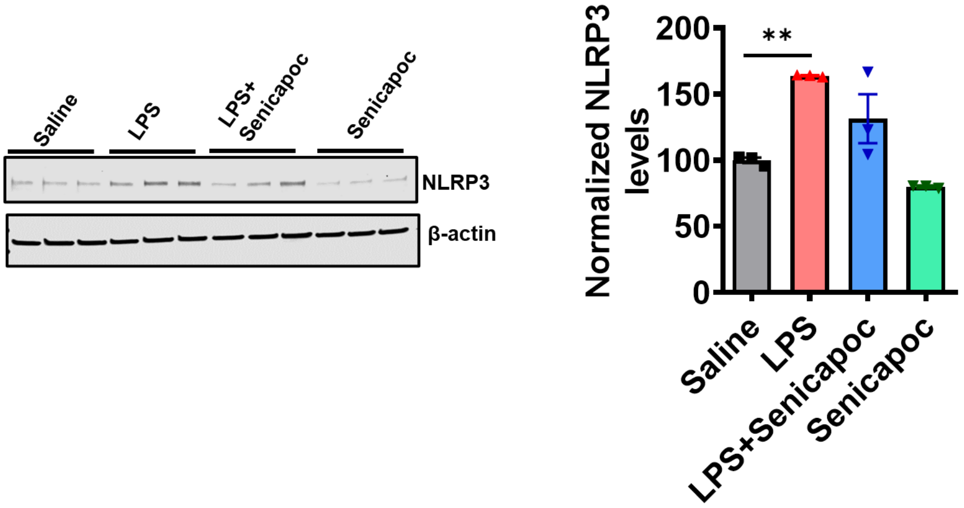


**Fig S4.** (A) Evaluation of cortical brain concentration of senicapoc using ultra-high-pressure liquid chromatography/mass spectrometry analysis reveals target engagement by the end of either 12 or 24 h post-drug administration. Senicapoc attains a concentration that is sufficient for KC3.1 blockade in the brain following i.p. administration of 50 mg/kg twice daily prior to LPS treatment followed by another dose on the day of LPS treatment. (B) Senicapoc pharmacological inhibition attenuates the NLRP3 expression induced by LPS in the mouse striatum. Data represented as ± SEM; n=4 mice per treatment group. *p≤0.05, **p<0.01.

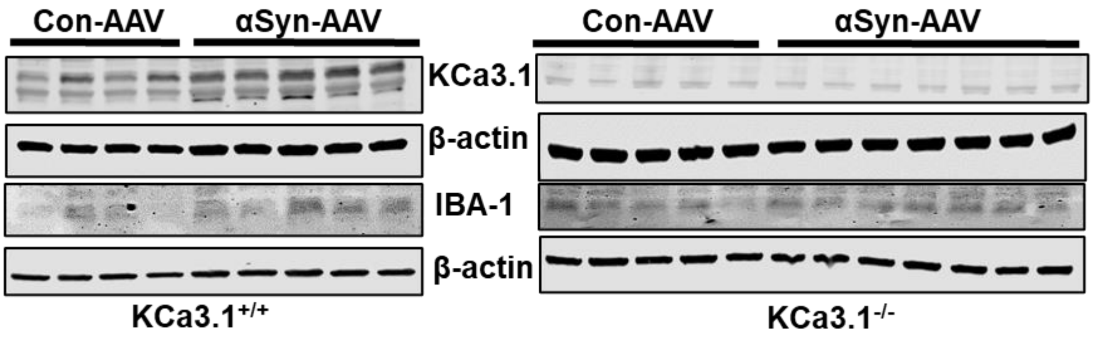


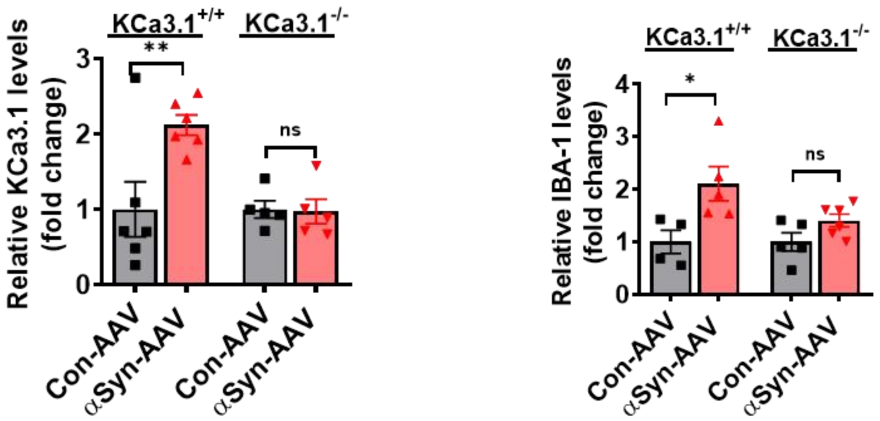


**Fig S5.** Global knockout of KCa3.1 reduces the IBA-1 expression in AAV-inoculated mice. Representative IBA-1 immunoblotting of striatal brain lysates of WT and KCa3.1 KO mice that received either control-AAV or αSyn-AAV.

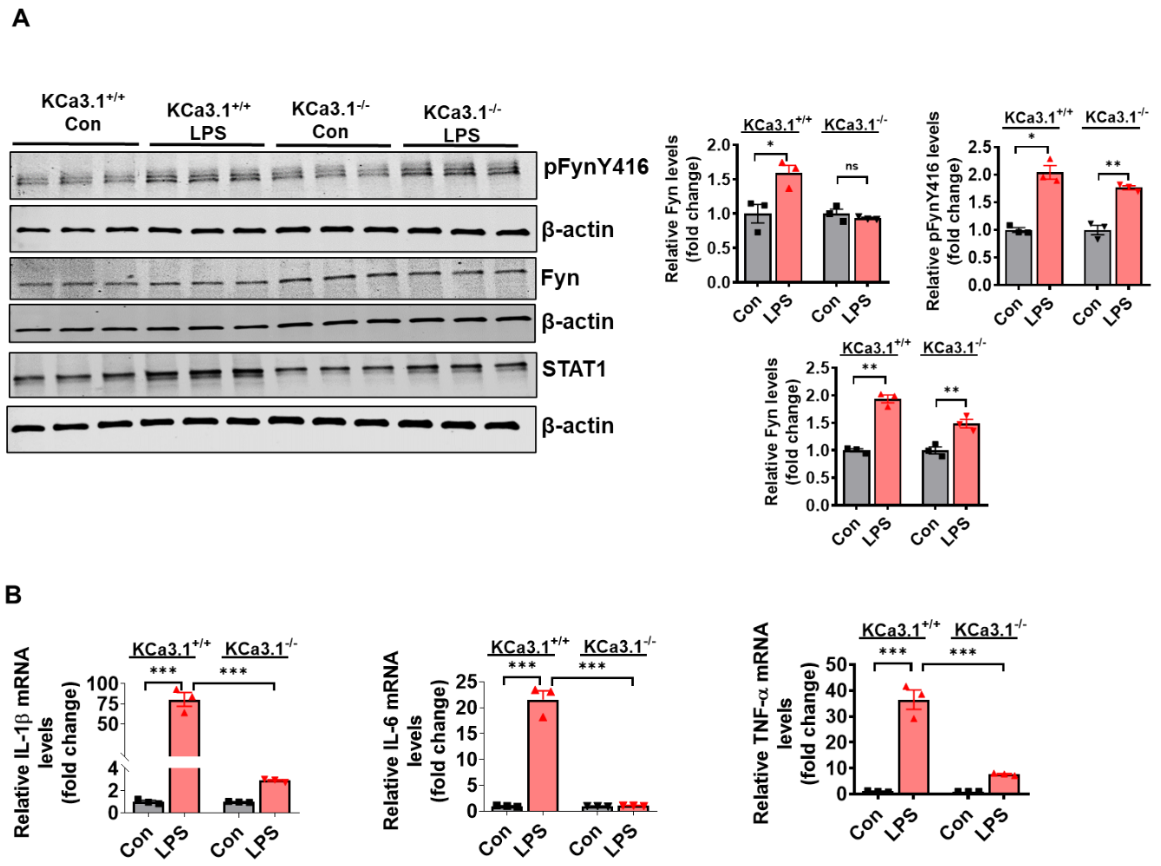


**Fig S6.** Genetic deletion of KCa3.1 alters Fyn-mediated STAT1 activation in microglia. (A) Immunoblots of Fyn, pFynY416, STAT1 in WT and KCa3.1 KO microglial cells following exposure to LPS (5 mg/kg for 18 h). (B) RT-qPCR analysis of TNF-α, IL-1β, IL-6 mRNA levels in adult microglia from control and KCa3.1 KO mice treated with or without LPS (5 mg/kg i.p. for 18 h). Data are presented as mean ± SEM; n=4-6 per group, Significance was determined by one-way ANOVA followed by Bonferroni test. *p≤0.05, **p<0.01, ***p<0.001.
